# Activation of the hypothalamic feeding centre upon visual prey detection

**DOI:** 10.1101/078527

**Authors:** Akira Muto, Pradeep Lal, Deepak Ailani, Gembu Abe, Mari Itoh, Koichi Kawakami

## Abstract

The visual system plays a major role in food/prey recognition in diurnal animals, and food intake is regulated by the hypothalamus. However, whether and how visual information about prey is conveyed to the hypothalamic feeding centre is largely unknown. Here we perform real-time imaging of neuronal activity in freely behaving or constrained zebrafish larvae and demonstrate that prey or prey-like visual stimuli activate the hypothalamic feeding centre. Furthermore, we identify prey detector neurons in the pretectal area that project to the hypothalamic feeding centre. Ablation of the pretectum completely abolishes prey capture behaviour and neurotoxin expression in the hypothalamic area also reduces feeding. Taken together, these results suggest that the pretecto-hypothalamic pathway plays a crucial role in conveying visual information to the feeding centre. Thus, this pathway possibly converts visual food detection into feeding motivation in zebrafish.

## Introduction

Control of feeding behaviour is one of the roles for the hypothalamus of the vertebrate brain. In mammals, the feeding centre resides in the lateral hypothalamus^1, 2^. In teleost fish, the inferior lobe of the hypothalamus (ILH) has been suggested to have crucial roles in feeding behaviour^3^. Ablation of these areas in the hypothalamus abolishes feeding behaviour^4^. Conversely, stimulation of these areas leads to feeding behaviour in both mammals and teleost fish^1, 5^. Thus, the important role of the hypothalamus in the regulation of feeding behaviours is highly conserved among vertebrates.

Neuronal activity in the feeding centre of the hypothalamus is affected not only by internal signals such as blood sugar levels, but also by the recognition of the availability of food sources in the environment. For example, neuronal activity has been observed in the hypothalamic feeding centre during appetitive and consummatory behaviours in rodents^6^ and at the sight of familiar food in monkeys^7^. However, because these studies used animals with prior feeding experience, it is not clear whether sensory-driven hypothalamic activity for appetitive behaviour is innate or learned through experience. In lower vertebrates that receive no parental care, larvae must attempt to catch prey, even at their first encounter, for their survival.

Here, we use zebrafish larvae as a vertebrate model to identify the neuronal pathways linking the visual prey perception to prey capture behaviour. Zebrafish start to swim at 4 day post-fertilization (dpf) and immediately start feeding. However, they can survive without feeding until 7 dpf because they can utilize their yolk as an energy source. Thus, the zebrafish is a good model system to study the neuronal activity in feeding behaviour at the earliest stage of life. In addition, the zebrafish is a genetically tractable animal model, and the transparent larvae brains enables the imaging study of functional neural circuits. In order to genetically label substructures in the brain or subpopulations of cells in specific neural circuits, we and other groups have established the use of the Gal4-UAS system in zebrafish^8, 9 10, 11, 12^ The activity of subsets of neurons that express Gal4 can be imaged with functional probes^13, 14^ or silenced with neurotoxins^8, 15^ to study the link between neuronal activity and behaviour.

Feeding in zebrafish larvae primarily depends on vision^16, 17^ The initial step in the visual recognition of possible prey or other small objects takes place in the retina in lower vertebrates^18, 19^, including zebrafish^20^. The optic tectum of the midbrain, which is the largest retinorecipient structure, is involved in the computation of object size^21, 22, 23, 24^ In addition, the optic tectum serves as a visuotopic map upon which the perceived prey is located^14^. Although direct projections from the visual system to the premotor reticulospinal neurons in the nucleus of the medial longitudinal fascicle (nMLF) have been demonstrated^20^, the neural pathway from these visual prey detection centres to the hypothalamic feeding centre is not known. We hypothesised that there should be a neural pathway that conveys visual information about prey to the hypothalamic feeding centre and that the presence of possible prey evokes hypothalamic activity that drives appetitive behaviour in zebrafish.

By imaging neuronal activity in free-swimming zebrafish larvae^25^, we investigate whether the visual detection of prey is sufficient for hypothalamic activation in naïve zebrafish larvae. Furthermore, we aim to identify the neural connections between the visual system and the hypothalamus. Here, we report our findings on the role of a pretectal nucleus as the prey detector and its anatomical and functional connection to the hypothalamic feeding centre in zebrafish larvae.

## Results

### The hypothalamus is activated at the sight of possible prey

In order to genetically define the postulated hypothalamic feeding centre, the inferior lobe of the hypothalamus (ILH) in zebrafish, we performed large-scale gene-/enhancer-trap screens^26^ and identified one enhancer trap line, hspGFFDMC76A, that expressed Gal4 in the ILH^27^ (Supplementary Fig. 1). In the adult zebrafish brain, UAS:EGFP reporter gene expression was observed throughout the entire structure of the ILH, including the dorsal zone of the periventricular nucleus, diffuse nucleus, central nucleus, and corpus mamillare. In addition, the reporter gene was also expressed in the torus lateralis and preoptic area (Supplementary Fig.1b-g). A similar pattern of expression, in addition to epithelial expression, was observed at the larval stage (Fig.1a). Although the subdivisions of the ILH in the larval brain have not yet been described, we observed at least two large populations of cells in the ILH at 5 dpf. The anterior population projected posteriorly and formed a tract that passed through the vagal lobe^28^ to the caudal end (Supplementary Fig. 1h). In addition to the ILH, two other populations of cells were labelled in the preoptic area both at the larval and adult stages (Fig. 1a, Supplementary Fig. 1d and h).

**Figure 1.**
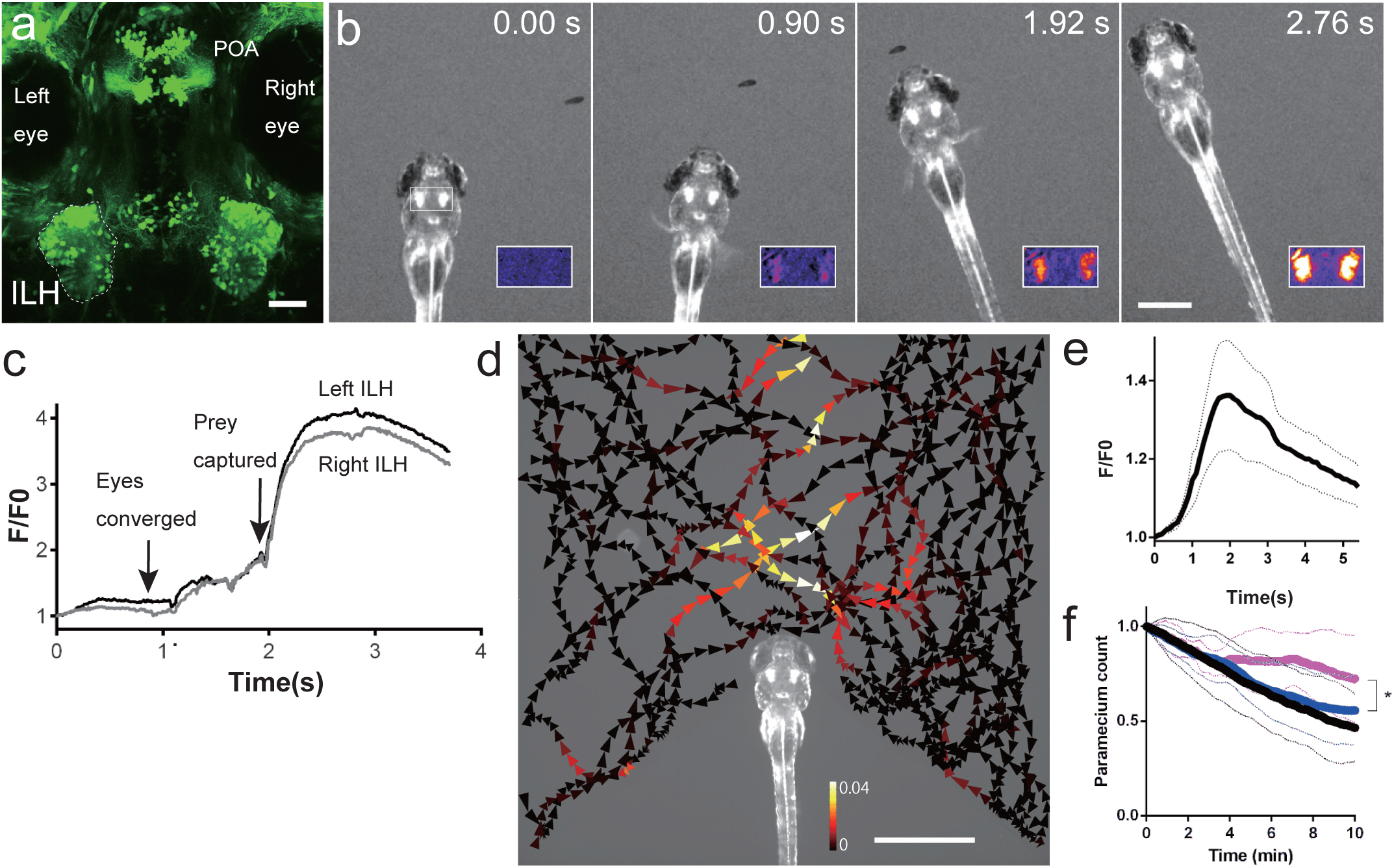
Activity in the ILH in the presence of prey. **a,** UAS:EGFP reporter gene expression in the ILH in hspGFFDMC76A Gal4 fish at 5 dpf. The left ILH is encircled by dotted lines. ILH: the inferior lobe of the hypothalamus, POA: preoptic area. Scale bar: 50 μm. **b,** ILH activation during prey capture behaviour in a previously unfed 4 dpf larva. Selected frames from a time-lapse fluorescent microscopy movie of UAShspzGCaMP6s;hspGFFDMC76A larvae. Inset: F/F0 in pseudo-colours. Scale bar: 500 pm. **c,** Normalised GCaMP6s fluorescence intensity (F/F0) from the recording shown in **b**. **d,** ILH activity in the presence of a paramecium in a 5 dpf zebrafish larva that was embedded in agarose. The trajectories of a single paramecium over 149 s are shown with the colour-coded changes in the intensity of the GCaMP6s fluorescence in the ILH. The signals in the left and right ILH were averaged. The length of the arrowhead indicates the distance travelled by the paramecium in 120 ms. Scale bar: 1 mm. **e,** ILH activity (GCaMP6s fluorescence intensity, F/F0) in response to a moving spot on a screen. (Mean (thick line) ± S.E.M (thin dotted lines), n = 5 larvae). The signals measured in the left and right ILH were averaged. A moving spot of 2.4° in diameter was presented to the larva at a speed of 100° /s from 0 to 3.1 s. **f,** Paramecium consumption in UAS:zBoTxBLCGFP;hspGFFDMC76A larvae (n = 6, magenta), control larvae (UAS:zBoTxBLCGFP alone) (n = 18, blue), and wild-type control larvae (n=13, black). The number of the paramecia left in the chamber was normalised to the initial count. The thick lines represent the mean and the thin dotted lines represent the standard deviation. The asterisk indicates a significant difference between the toxin:Gal4 double transgenic larvae and UAS:zBoTxBLCGFP alone control larvae at 10 min (one-tailed t-test, p=0.040).

To visualise neuronal activity in this hypothalamic feeding centre, we generated UAShspzGCaMP6s^29^ transgenic zebrafish and expressed the calcium (Ca) probe GCaMP6s in the hspGFFDMC76A Gal4 line. We performed calcium imaging in previously unfed zebrafish larvae to determine whether the ILH was activated during prey capture behaviour. GCaMP6s fluorescence intensity was increased in the ILH when a paramecium approached the larva (just prior to eye convergence, F/F0 =1.3 ± 0.12 (mean ± S.D.), n = 24 events in 4 larvae), just prior to the rapid capturing movement (Peak F/F0 = 1.7 ± 0.28, n = 8 events in 4 larvae) and after completion of prey capture (Peak F/F0 = 3.46 ± 0.40, mean ± S.D., n = 4 larvae; Fig. 1b-c, Supplementary Fig.2, Supplementary Movie 1). Although the ILH showed bilateral activation, the ILH on the opposite side of the paramecium location showed higher elevation of the calcium signal at the time of eye convergence (Supplementary Fig.2, in 19 cases out of 22 in 4 larvae, p=0.003, binominal test).

Since image registration could not completely eliminate movement artefacts in the quantification of the calcium signals, we performed control experiments with UAS:EGFP-expressing larvae to assess the extent of the movement artefacts. In our experimental conditions, the fluctuations in the intensity of the EGFP fluorescence in the ILH after image registration were negligible, which suggested that the changes in the GCaMP6s fluorescence reflected actual calcium signals (Supplementary Fig.3). Next, to examine the optimal positions of the prey in the visual field for the ILH responses, a larva was embedded in agarose, and a single paramecium was presented in the recording chamber. The presence of a paramecium in front of the larva evoked a response in the ILH, regardless of the movement direction of the prey (Fig. 1d, Supplementary Fig. 4, Supplementary Movie 2).

Living paramecia may contain olfactory cues as well as visual cues. To examine whether a visual cue was sufficient to evoke this hypothalamic activity, we showed a moving spot, which mimicked the paramecium, to a larva that was embedded in agarose while the brain was imaged with a confocal microscope. The moving spot evoked the ILH responses that were comparable in amplitude to the calcium signals evoked by living prey (Fig. 1e), which suggested that vision is the major sensory modality responsible for the activation of the ILH in prey capture behaviour.

To test the necessity of the neuronal activity in the ILH for the prey capture behaviour, we crossed this ILH Gal4 line with a botulinum toxin line, UAS:zBoTxBLCGFP and then assessed the prey capturing ability of the larvae. We found that paramecium consumption was significantly decreased in the larvae that expressed botulinum toxin in the ILH (Fig. 1f). Expression of the toxin did not affect the locomotor activity (Supplementary Fig. 5a), and larvae carrying UAS:zBoTxBLCGFP alone or with another gal4 SAGFF(LF)27A^30^ did not show significant change in prey capture activity (Supplementary Fig. 5c). These results suggested that the hypothalamic activity is important for the feeding behaviour. We did not observe a significant reduction in eye convergence in the larvae that expressed the toxin in the ILH (Supplementary Fig. 5b).

### Neurons in the pretectum serve as prey detectors

In order to investigate the neuronal pathway that links visual prey recognition and hypothalamic feeding regulation, we next tried to identify the neuronal module in the brain that responded to prey, other than the optic tectum of the midbrain that shows visuotopic activity^14^. In our collection of Gal4 gene-/enhancer-trap lines, we expressed GCaMP6s in various regions in the brain and then imaged the larval brains while the larvae were in the presence of paramecia. This approach revealed that cells in the pretectal area in the gSAzGFFM119B line (hereafter, referred to as the 119B-pretectal cells) showed calcium signals that correlated with the proximal presence of a paramecium (Fig. 2a-c, Supplementary Fig. 6, Supplementary Movie 3). In addition to the pretectal area, gSAzGFFM119B expression was observed in the dorsal half of the forebrain, olfactory bulb, preglomerular nuclei, optic tectum, and cerebellum. In contrast to the pretectal cells, none of these areas showed apparent calcium signals at the sight of prey. Since moving artefacts are inevitable when imaging of free-swimming larvae, we performed control EGFP recordings. Smaller regions of interest (ROIs) in the pretectum generated much larger artefacts in the traces of fluorescence intensity than those observed in the ILH after the larvae started to move to capture the prey. Even in this condition, the EGFP fluorescence did not increase before eye convergence (Supplementary Fig.7). These control experiments validated the GCaMP6s signals in a condition involving minimal movements of the larvae. To characterize the activity of the 119B-pretectal cells, we again used the experimental set up of the agarose-embedded larvae. Visually responsive neurons in the pretectal area were not selective for the swimming direction of the paramecium and their responses were not necessarily associated with behaviour (e.g. eye convergence^17^), which indicated that this pretectal neuronal activity was involved in the visual detection of prey (Fig. 2d, Supplementary Fig. 8, Supplementary Movies 4-5).

**Figure 2.**
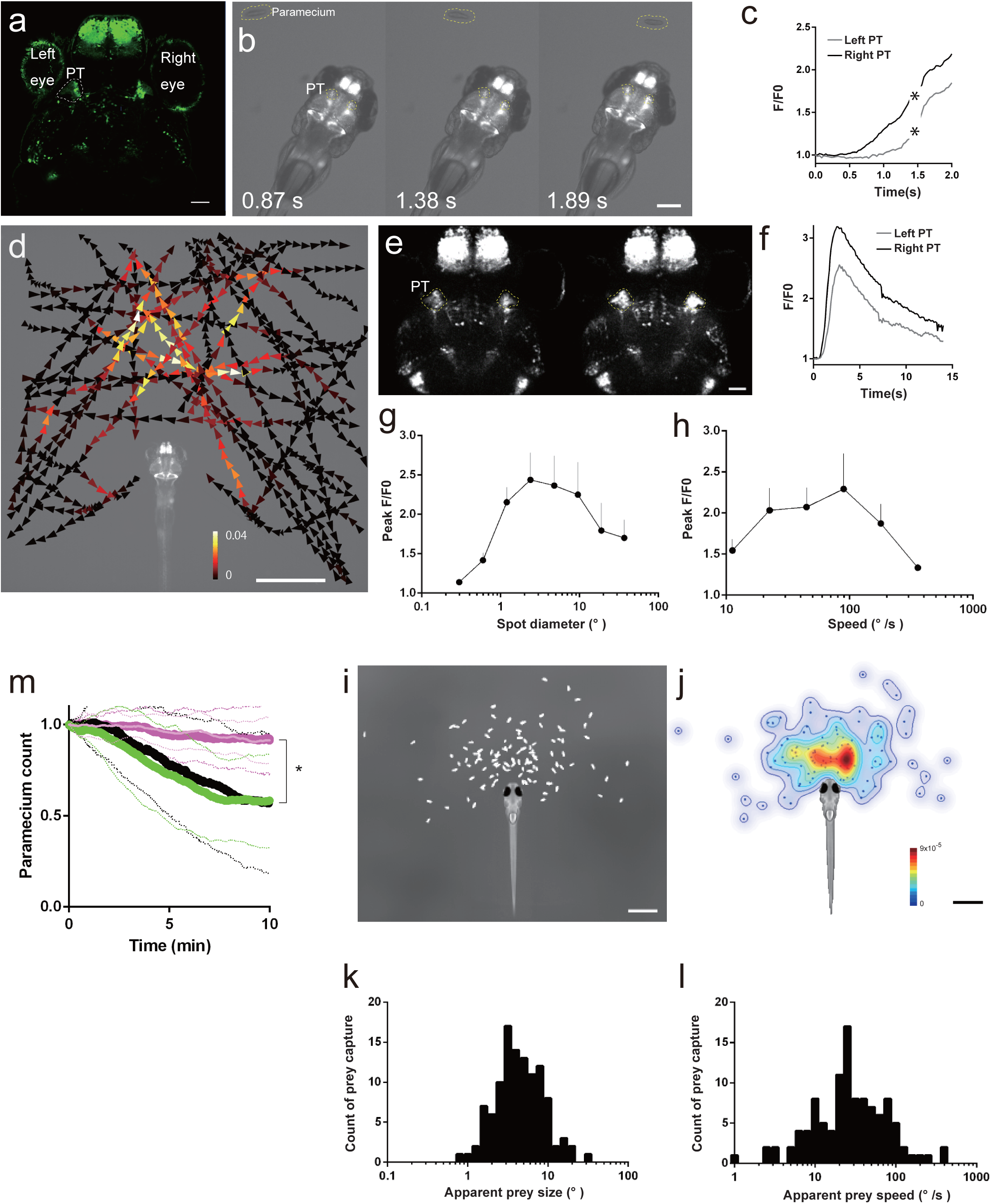
Activity in a subset of cells in the pretectal area in the presence of prey. **a,** UAS:EGFP reporter gene expression in the pretectal area (PT, encircled in dotted lines) in gSAzGFFM119B Gal4 larva at 5 dpf. Scale bar: 50 pm. **b,** Pretectal activation in response to prey at 4 dpf. GCaMP6s fluorescence images. Scale bar: 500 pm. **c,** GCaMP6s fluorescence intensity in the pretectal areas shown in **b**. Asterisks: missed data points due to large movements of the larva. **d,** Averaged activity of the bilateral pretectal areas in the presence of prey in a 6 dpf larva embedded in agarose. The trajectories of a single paramecium over 107 s with colour-coded changes in the intensity of GCaMP6s fluorescence in the pretectal areas. Scale bar: 1 mm. **e,** Confocal images showing pretectal activation in response to a moving spot. Scale bar: 50 μm. **f,** GCaMP6s fluorescence intensity obtained from the time-lapse images in **e**. A moving spot of 2.4° in diameter was presented to a larva at a speed of 100° /s from 0.3 to 3.4 s. **g** and **h,** Size-tuning and speed-tuning curve, respectively (n = 6 larvae at 4 dpf; Mean ± S.E.M.). Maximal responses to a moving spot. The x-axis is logarithmically scaled. **i,** Relative positions of the paramecium at the onset of prey capture in free-swimming larvae at 5 dpf (n = 110 successfully completed prey capture events by 6 larvae). Scale bar: 1 mm. **j,** Probability density of the paramecium positions shown in **i**. Colour bar:0 (blue) and 9x10^-5^ (red). Scale bar: 1 mm. **k** and **1,** Histograms of the apparent sizes and speeds, respectively, of the prey at the onset of prey capture from the data shown in **i**. **m,** Paramecium consumption in the UAS:EGFP;gSAzGFFM119B larvae that were subjected to bilateral pretectal ablation (n = 10, thick magenta line) or unilateral (n=10, thin light-magenta line), UAS:EGFP;gSAzGFFM119B larvae subjected to bilateral olfactory bulb ablation (n = 11, green), and UAS:EGFP;gSAzGFFM119B larvae (no laser irradiation, n=8, black). Solid lines:mean. Dotted lines:S.D. Asterisks: significant differences between each of pretectum ablation groups and each of control groups at 10 min (Tukey’s HSD test, p=0.0016 for bilateral pretectum ablation and olfactory bulb ablation).

In order to test if the observed pretectal responses can be evoked solely by visual stimuli, we showed a moving spot on a small screen to an agarose-embedded larva. Both the left and right pretectum showed activation in response to the visual stimulus (Fig. 2e-f, Supplementary Movie 6). A moving spot with the size of 2.4° in diameter and at a speed of 100° /s evoked the strongest response (Fig. 2g-h). To investigate whether these size-tuning and speed-tuning of the pretectal responsiveness were related to prey capture behaviour, we next observed natural prey capture behaviour and measured the apparent size and speed of the prey at the onset of prey capture behaviour. In the majority of the prey capture events, the apparent prey size and speed were comparable to the size tuning and speed tuning of the pretectal responsiveness, respectively (Fig.2i-l).

Next, we tested the necessity of the 119B-pretectal cells during prey capture behaviour. We ablated these cells with a two-photon laser at 4 dpf and tested the prey capture ability at 5 dpf (Supplementary Fig. 9a-c). Both bilateral and unilateral ablations abolished prey capture behaviour (Fig. 2m, Supplementary Movie 7), which suggested that bilateral activity is necessary for prey capture. The rate of occurrence of eye convergence was reduced, whereas the optokinetic response, which is another visuomotor behaviour, remained unaffected (Supplementary Fig. 9d-e). Locomotor activity was not affected by laser ablation of the pretectum (Supplementary Fig.9f). As a control experiment for laser ablation, we ablated a subpopulation of the olfactory bulb cells that were labelled in gSAzGFFM119B, whose number was comparable to that of the 119B-pretectal cells (Supplementary Fig.9c). The olfactory bulb-ablated larvae showed prey capture activity that was similar to untreated control larvae (Fig. 2m).

### The prey detector neurons connect to the hypothalamus

In order to identify the efferents of the 119B-pretectal cells, we sparsely labelled Gal4-expressing cells by injecting the UAS:EGFP reporter DNA into the eggs of the gSAzGFFM119B Gal4 line. We observed that some of the 119B-pretectal cells projected to the caudomedial part of the ILH^31^ (Fig. 3a, Supplementary Fig.10, Supplementary Movie 8). In the UAShspzGCaMP6s; gSAzGFFM119B;hspGFFDMC76A double Gal4 transgenic larvae, the activities of the pretectal area and the ILH were correlated significantly (correlation coefficient > 0.8) (Fig. 3b, Supplementary Movie 9). Unilateral ablation of the 119B-pretectal cells revealed that both the 119B-pretectal cells and the ILH on the unablated side were more responsive to the prey when it was present in the other hemifield (Fig. 3c-e, “Contralateral pretectum” and “Contralateral ILH”, Supplementary Fig.11). Residual activity were present in the ablated pretectum probably due to the remaining cells that survived laser-ablation. Bilateral ablation of the pretectum dramatically reduced activity in the ILH (Fig.3e, Supplementary Fig.12). These results suggest a functional connection between the 119B-pretectal cells and the ILH, although the connectivity may not be necessarily all direct, but it could be indirect involving other relay areas.

**Figure 3.**
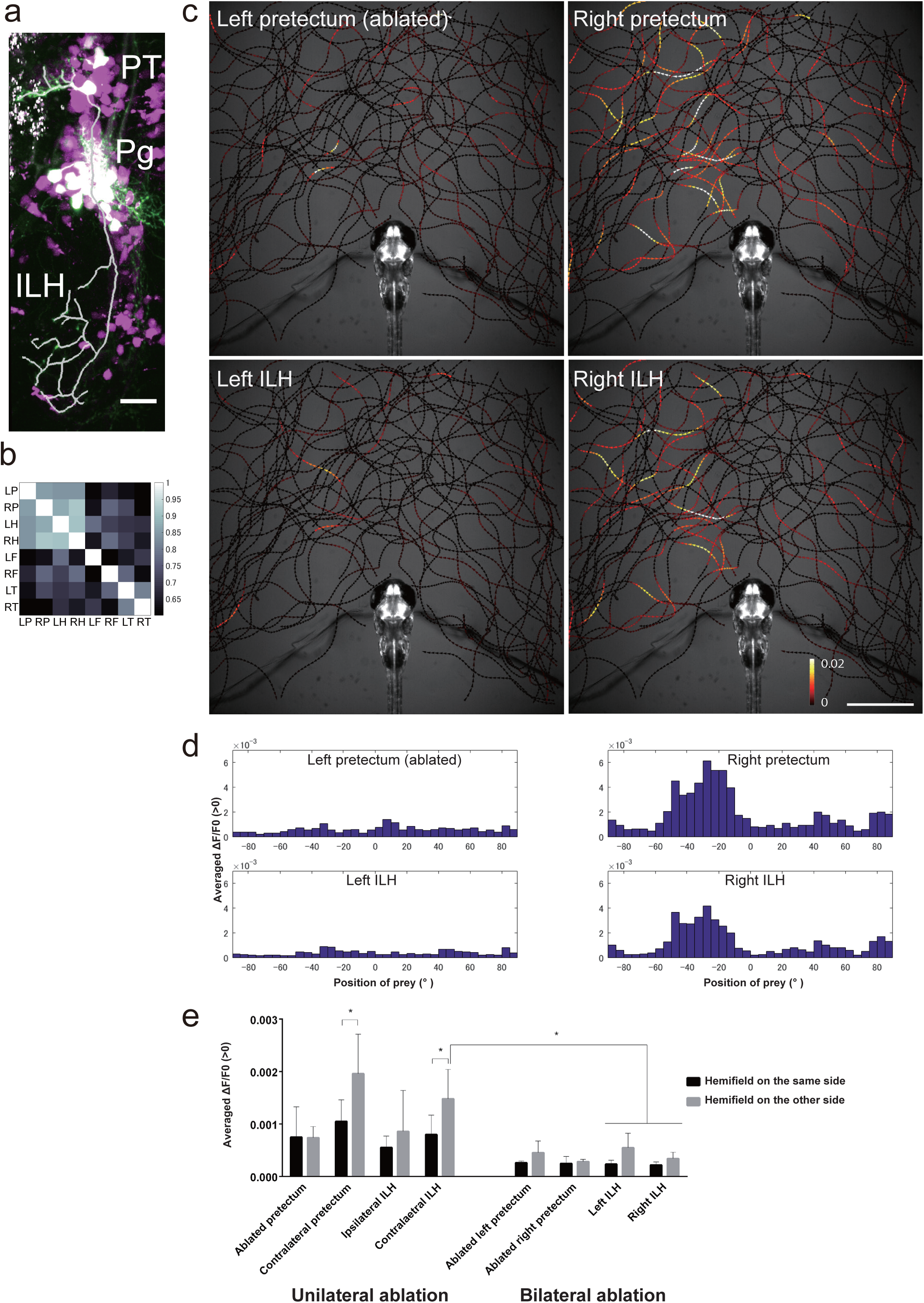
Anatomical and functional connections from the pretectum to the ILH. **a,** Merged image of the pretectal cells that were sparsely labelled by the injection of UAS:EGFP DNA (EGFP in green) in the UAS:RFP;gSAzGFFM119B background (red fluorescent protein shown in magenta). The axonal projection of a single pretectal cell is shown as a thick white line. ILH, the inferior lobe of the hypothalamus; PT, pretectum; Pg, Preglomerular nuclei. Scale bar: 20 μm. **b,** Correlation of the activity of the pretectum and ILH. UAShspzGCaMP6s;hspGFFDMC76A;gSAzGFFM119B double Gal4 transgenic larvae were imaged and the cross-correlation coefficients of the calcium signals in the specified areas were calculated (Colour-coded, 1: highest correlation, 0: no correlation). LP, left pretectum; RP, right pretectum; LH, posterior half of the left hypothalamus (ILH); RH, posterior half of the right hypothalamus (ILH); LF, left forebrain; RF, right forebrain; LT, left optic tectum; RT, right optic tectum. **c,** Activities in the pretectum and ILH in the presence of a paramecium in a 5 dpf larva that was embedded in agarose. The trajectories of a single paramecium over 318 s are shown with colour-coded changes in the intensity of GCaMP6s fluorescence in a 5 dpf UAShspzGCaMP6s; hspGFFDMC76A;gSAzGFFM119B double Gal4-transgenic larva that was subjected to laser ablation of the left pretectum. Scale bar: 1 mm. **d,** Preference for the azimuthal position of the paramecium. **e,** Left-right hemifield preference in the pretectum and ILH in 5 dpf larvae that were subjected to unilateral pretectum ablation (bar graphs on the left) and abolition of the neuronal activity by bilateral pretectum ablation (bar graphs on the right). Mean ± S.D. The asterisks indicate significant differences. “Contralateral pretectum”: unilaterally ablated larvae n=6, two-tailed t-test, p=0.0261. “Contralateral ILH”: unilaterally ablated larvae n=6, two-tailed t-test, p=0.0325. “Contralateral ILH” - “Left ILH” and “Right ILH”: bilaterally ablated larvae n=4, Tukey’s HSD test, less than p<0.0034 for all 4 groups. On the x-axis are the brain areas on which regions of interest (ROIs) were set to measure the GCaMP6s fluorescence intensities.

### Locations of the 119B-pretectal area and RGC arbours

In zebrafish, retinal ganglion cells (RGCs) project to the optic tectum and several nuclei in the pretectal area, where they form characteristic arborisation fields (AFs)^32, 33^. The optic tectum (AF-10) is the largest retinal target and other smaller distinct areas are named AF-1 to AF-9^32^. To examine whether the 119B-pretectal cells received a direct input from the retina in larval zebrafish, we injected a lipophilic dye, DiI, into an eye to label the axons of the RGCs (Supplementary Fig. 13). The 119B-pretectal cells were found to be located ventral to the AF-7 area. Some bundles of the optic tract passed through the pretectal neuropil area; however, they did not arbourise in the dendritic area of the 119-pretectal cells (Supplementary Fig. 13d). No overlap of the 119B-pretectal area with any of the major AFs suggests that the 119B-pretectal cells are not a major direct target of the RGCs.

## Discussion

In this study, we observed neuronal activity in the inferior lobe of the hypothalamus (ILH) that was driven by the visual prey perception in zebrafish larvae without any prior feeding experience. We also identified prey detector neurons in the pretectal area that projected directly to the ILH and were indispensable for prey capture. We propose that visual information conveyed by this pretecto-hypothalamic circuit is essential for modulatory function of the hypothalamus in feeding behaviour (Supplementary Fig. 14).

The ILH is a conspicuous structure bulging out on the ventral brain surface in adult zebrafish brain but at larval stage it has not been unequivocally identified due to the lack of a reliable marker. Taking advantage of our collection of the gal4 gene-/enhancer-trap lines^26^, we used hspGFFDMC76A Gal4 line to label the ILH, which showed subdivision into at least two areas, anterior and posterior, at larval stage. The cells in the anterior part of the ILH formed an outgoing tract that passed along the vagal area. The two tracts on the left and right sides converged at the posterior end of the vagal area. In goldfish and tilapia, the central nucleus and the nucleus of lateral recess of the ILH project to the vagal lobe and the commissural nucleus of Cajal^34, 35^. Recent work on adult zebrafish also showed that the ILH forms a part of the gustatory circuit^36^. Our observations in larval zebrafish are consistent with those studies and demonstrate that these anatomical structures are conserved among these species. The posterior part of the ILH received gSAzGFFM119B Gal4-labeled pretectal efferents. Based on this input pattern and the expression in the adult brain, we identified this area as the corpus mamillare.

The ILH has been hypothesized to be the feeding centre, or “hypothalamic feeding area”^3^, in teleost fish based on surgical lesion^4^ and electrical stimulation^5^ experiments. As the ILH has a complex structure with subdivisions and intricate fibre connections, perturbations of the ILH function by these classical experimental tools could not be very specific to the targeted areas. More specific interventions are required to determine exact roles for the ILH and its subdivisions. Our observation of neuronal activity in the ILH during initial prey perception and succeeding prey capture behaviour supports the notion that the feeding centre in teleosts resides in the ILH. In the present study, we only used previously unfed larvae at 4-6 dpf, which probably corresponds to a hungry condition as zebrafish larvae start foraging at 4 dpf. If the ILH activity represents feeding motivation, it would be lowered by feeding. A study in monkeys indeed showed that satiety reduces hypothalamic visual response to the food^37, 38^. In zebrafish, the feeding state was shown to modulate visual perception in the optic tectum by shifting the size-tuning property through a serotonergic system^22^. The pretecto-hypothalamic circuit might serve as another target for the modulation of this visually driven behaviour by hunger/satiety state.

Prey detector is a long standing notion in neuroethology, and its neural correlate has been a topic of great interest^39^. Studies in zebrafish and other species have suggested that prey detection is initiated in the retina and further elaborated in the optic tectum for the extraction of the visual features such as size and motion of the prey^21, 22, 23, 24, 40, 41^. Being either retinorecipient or tectorecipient^27^, pretectal nuclei also participate in the neural circuit for prey perception. In zebrafish, the RGCs project to the optic tectum and also to distinct pretectal areas where they form arbourisation fields (AFs)^32, 33^. A subpopulation of the RGCs that project to the AF-7 pretectal area are involved in prey capture^20^. The pretectal area containing the cells that receive inputs from AF-7 was identified as the parvocellular superficial pretectal nucleus (PSp), and these cells project to the nMLF, motor area in the hindbrain, and also to the optic tectum in zebrafish^20^. In a perciform teleost, the retinorecipient cells in the pretectal nucleus corticalis (NC), which is absent in zebrafish^42^, show visual responses to a small moving spot and send projections to the optic tectum. The receptive fields of these NC neurons are large and cover the entire visual field^43^. In contrast with the results of these previous studies, the 119B-pretectal cells seem not to receive direct inputs from the RGCs, respond to an object located in a relatively limited area in front of the larva, and project to the caudal part of the ILH (the corpus mamillare). These differences suggest that 119B-pretectal cells are different cell populations from previously reported cell types that respond to small objects. In addition to the ILH projection, we observed another projection from the 119B-pretectal cells toward the precerebellar area, possibly to the nucleus lateralis valvulae^44^ (Supplementary Movie 10). Based on these morphological and functional observations, the 119B-pretectal cell group was identified as the magnocellular superficial pretectal nucleus (PSm)^42^ Thus, our study along with Semmelhack’s^20^ suggest that two distinct pretectal nuclei, namely, PSp and PSm, are involved in prey perception in zebrafish. PSm-pathway might have modulatory role over the more directly motor-related Psp-pathway. The significance of having a dual system of prey perception and its interrelationships remain to be elucidated.

In neuroethology, prey-catching behaviour has been extensively studied in toads. Thalamic-pretectal lesions in toads disinhibit the size-specificity of prey-catching behaviour, which results in enhanced prey-catching behaviour to larger stimuli that normally cause an avoidance response^45^. Thus, the pretectum in toads appears to function in a manner opposite to that observed in our study. This discrepancy might be due to differences in the ablated areas or to evolutionarily divergent functions of the pretectal nuclei^27^.

In our calcium imaging in free-swimming zebrafish larvae, the elevation of the ILH activity started before the onset of behavioural response (i.e., eye convergence). Although this suggests correlation of the ILH activity with the initiation of the prey capture behaviour, we did not detect a significant reduction of the eye convergence time in the larvae with botulinum toxin expression in the ILH. This could be partly due to an insufficient expression level of the botulinum toxin in the ILH. Alternatively, other downstream pathways might be more crucial for the initiation of prey capture behaviour and the ILH silencing had an effect after prey capture was started. Further study is necessary to clarify the causality between the ILH activity and behavioural processes.

Both in the 119B-pretectal cells and the ILH, bilateral activity was evoked merely by the presence of prey and the activity was not necessarily associated with eye convergence (the initial step in the prey capture behaviour). In the cases where eye convergence occurred, the intensity of the GCaMP6s fluorescence was observed to increase before the eye movements. This suggests that the activation of the pretecto-hypothalmic circuit, at least in the early phase, is solely evoked by visual prey detection. However, it is notable that larger calcium signals in the 119B-pretectal cells were often observed in the cases with the eye convergence. The119B-pretectal cells also projected to the precerebellar area which possibly relays information to the cerebellum. Cerebellar activities were observed to be associated with eye movements. Existence of diverging neural pathways and their activity suggests more roles for the 119B-pretectal cells than initial perception of the prey.

Prey capture circuit in zebrafish is a good model system to study goal-directed sensorimotor transformation and its modulation by internal states, external conditions and feeding experience. To fully understand this behaviour, we need to elucidate the entire circuit. Since the PSm receives input from the optic tectum in carps (a species that is closely related to zebrafish), the 119B-pretectal cells might also receive input from the optic tectum^44^. While progress has been made on the visuomotor circuit for prey capture, it is still largely unknown how gustatory information, that usually comes seconds after initial prey detection, can modulate the visual prey perception, a critical pathway for the reinforcement learning of food/non-food discrimination in a given feeding environment. In the ILH, much larger calcium signals were observed after the completion of prey capture compared with visually evoked calcium signals. This massive neuronal activity in the late phase of the behaviour might be associated with gustatory sensations of the prey and/or chewing or swallowing. The observation of a sustained activity in the ILH from the initial prey perception to completion of prey consumption, and the fact that the ILH receives visual input and gustatory feedback tempt us to speculate that the ILH might serve as the place for integration of visual and gustatory information of prey. The multimodal integration should be implicated in reinforcement of the behavioural decision based on the visual features of prey.

In this study, we demonstrated hypothalamic activation upon visual prey detection and genetically identified prey detector neurons in the pretectal area that had direct projections to this hypothalamic feeding centre. Since the visual recognition of familiar food elicits neuronal responses in the lateral hypothalamus in primates^7^, visually driven hypothalamic activation seems to be a fundamental process in vertebrate feeding behaviour. This innate motivational system is crucial for the survival of larvae that receive no parental care.

## Methods

### Zebrafish husbandry

Adult zebrafish were maintained at 25°C under a regular 13 h light/11 h dark cycle. Embryos were kept at 28.5°C under the same light/dark cycle until the larval stage.

### Generation of transgenic zebrafish

UAShspzGCaMP6s13A transgenic zebrafish were generated by injecting a DNA construct into fertilised eggs at the 1-cell stage. The DNA construct consisted of 5xUAS (the Gal4 binding site), the heat shock protein 70 promoter (650 bp), a zebrafish codon-optimised GCaMP6s (https://www.addgene.org/40753/), and a poly-A addition signal in a Tol2 transposon vector. GCaMP6s fish with a single insertion was identified by Southern blot analysis and used in this study. Generation of the Gal4 gene-trap lines, gSAIzGFFM119B and hspGFFDMC76A, are described elsewhere (manuscript under preparation). UAS:zBoTxBLCGFP42A transgenic zebrafish were generated by injecting a botulinum toxin light chain B DNA construct^15, 46^. Fish in F1 and F2 generation of the transgenic line were subjected to Southern blot analysis to isolate fish that harboured a single insertion of the UAS:zBoTxBLCGFP transgene. F2 fish were mated with the Gal4 lines and the larvae in F3 generation were used in this study.

### Identification of the loci of Gal4 insertions

Inverse PCR was performed and a fragment of the neighbouring genomic DNA was cloned and sequenced^47^ The nucleotide sequence was compared to the zebrafish Ensemble genome database (http://www.ensembl.org/Danio_rerio/Info/Index) to determine the insertion sites in the genome.

### Preparation of larvae for imaging and behavioural studies

Previously unfed 4–7 dpf larvae were used throughout the experiments. GCaMP6s was expressed in the larval brain by mating a Gal4 zebrafish with a UAShspzGCaMP6s transgenic zebrafish. To attain the transparency necessary for optical imaging, both Gal4 and UAS lines were maintained on a *nacre* background to produce *nacre* homozygotes that lacked melanophores (and hence, black pigment on the surface of the brain and body except in the retinal pigment epithelium)^48^.

### Calcium imaging with an epi-fluorescence microscope

For the imaging of a free-swimming larva, a single larva and a few paramecia were put in a small chamber (Secure-Seal Hybridization Chamber Gasket, 8 chambers, 9 mm diameter x 0.8 mm depth, Catalogue # S-24732, Molecular Probes, *Eugene,* OR, USA) and observed under an epi-fluorescence microscope (Imager.Z1, Carl Zeiss, Germany) with an objective lens (2.5X/NA0.12 or 5X/NA0.15) equipped with a scientific CMOS camera (ORCA-Flash4.0, model:C11440-22CU, Hamamatsu Photonics, Hamamatsu, Japan). Images were recorded at 100 fps (10 ms exposure). The XY-stage was manually moved to locate the larva at the centre of the camera view each time when it moved out of the camera view. Fluorescent images were acquired with time-lapse recording software (HCImage with a Hard Disk Recording module, Hamamatsu Photonics, Japan).

For the analysis of the fluorescent signals in free-swimming larvae, image registration was performed in two steps. In the first step, using LabVIEW’s template matching function in NI VISION module (National Instruments, Austin, TX, USA), the location (x,y) and the orientation (angle) of the larva in each frame were obtained.

Using these parameters, each frame was translated and rotated to place the larva in the same position through frames. Because the deviation of the larval position was not small enough, in the second step we performed image registration again using TurboReg plugin (http://bigwww.epfl.ch/thevenaz/turboreg/) in ImageJ (Rasband, W.S., National Institutes of Health, Bethesda, Maryland, USA, http://imagei.nih.gov/ii/).

For the imaging of an agarose-embedded larva, a single larva was mounted in 2% low melting agarose. Agarose around the head was removed to expose the eyes and to make space for paramecia to swim. A single paramecium was put in the chamber. Time-lapse recording was performed with an acquisition rate of 33.33 fps using the same microscope as mentioned above.

ImageJ was used to obtain mean pixel values (i.e. calcium signals) in a region of interest (ROI) set on a brain structure in the time-lapse movie. To read the .cxd format image files of ORCA-Flash4.0 in ImageJ, the Bio-Formats plugin was used^49^

Calcium signals were presented in either way, normalized GCaMP6s fluorescence intensity (F/F0) or increment of the GCaMP6s fluorescence intensity (“Averaged Ca rises”). In F/F0 representation, GCaMP6s fluorescence intensities (mean pixel values in a ROI set on the pretectum, the inferior lobe of the hypothalamus, or other neuronal structures) divided by F0 (fluorescence intensity at basal level). The signals were averaged for 11 time points (moving average) before the normalization.

The timings of the presented calcium signals in the main text are as follows: “Just prior to eye convergence”: 1 frame (=10ms) before noticeable inward movement of the eyes. “Just prior to a rapid capturing movement”: 1 frame (=10ms) before noticeable rushing movement.

For the “Averaged ∆F/F0 (>0)”, the changes in fluorescence intensity in inter-frame interval (30 ms) was calculated and averaged for 11 time points (moving average). Because of the slow decay of GCaMP6s signals^29^, which occurres in seconds, only the timing of the increased GCaMP6s signals is correlated with the timing of the stimulus presentation (i.e., approach of the paramecium). For this reason, only positive values (∆F>0) were held and negative values (∆F<0) were replaced with zero, in order to examine the presence of the correlated neuronal activity with the location of the swimming paramecium. Changes in the fluorescence intensity per frame were depicted on a colour-map (black: change ≤ 0; dark red - yellowish - white: change≥ 0). To generate a Ca signal map on the paramecium trajectories, a custom-made Matlab (Mathworks, Natick, MA, USA) script was used. To calculate orientation-preference, the position of the paramecium was measured by the angle (the midpoint of the two eyes of the zebrafish larva as the origin, upward: 0°, and clockwise positive), and all the increments (only positive values) of the GCaMP6s signal at each paramecium position within each 5° bin were averaged. In the analysis of laterality (i.e., preferred neuronal response to either left or right hemifield), the area below the level of ears in the image were excluded from the calculation of the averaged ∆F/F0 because prey in the area did not evoke neuronal response or prey capture behaviour.

To observe the correlated activity of the pretectum and the ILH, UAShspzGCaMP6s;gSAIzGFFM119B;hspGFFDMC76A larvae were used. In this case, the ROI for the ILH was set only in the posterior half so that it will not overlap with the optic tectum. ROIs for the optic tectum were set so as not to include the ILH area. Correlation of the activities in different neuronal structures was assessed by calculating cross-correlation coefficients using Matlab functions.

### Calcium imaging with a confocal microscope

The confocal laser scanning microscope system used for calcium imaging consisted of a microscope (Examiner, Carl Zeiss, Germany), a scanning unit with a Nipkow spinning disk (CSU-W1, Yokogawa Electric, Tokyo, Japan), and an EM-CCD camera (iXon DU-888, Andor, Belfast, UK). The imaging system was controlled by iQ3 software (Andor, Belfast, UK). A water immersion objective lens (W Plan-Apochromat 20X/1.0, Carl Zeiss, Germany) was used.

Transgenic zebrafish larvae carrying the UAShspzGCaMP6s;gSAIzGFFM119B or UAShspzGCaMP6s; hspGFFDMC76A on a *nacre* background were placed in an acrylic box (60 mm × 60 mm; 35 mm in height) filled with system water. One wall of the box was covered with a rear projection screen film (Type: GSK, Kimoto Co. Ltd, Saitama, Japan). The larvae were placed facing toward the screen at a distance of 20 mm and at the middle height of the box. Animations of a moving spot with different sizes and speeds were generated in MATLAB and projected onto the screen by a small projector (M115HD, DELL, TX, USA). A long-pass filter (FGL610, Thorlabs, Newton, NJ, USA) and a plano-convex lens (Thorlabs, NJ, USA) were set in front of the projector lens to remove the green component of the light and to enable focusing at a shorter distance, respectively. The timings of the start of calcium imaging and stimulus presentation were controlled by externally triggering the image acquisition software iQ3. Triggering signals were sent through a data acquisition module (NI USB-6211, National Instruments, controlled in a Matlab script for visual stimulus presentation), and their timings and shutter timing signals from the iXon DU-888 were recorded with another data acquisition module (NI USB-6008). Imaging of the larval brains was performed with a confocal microscope, as described above.

### Behavioural assay for prey capture

One zebrafish larva was placed in a recording chamber (diameter: 20 mm; depth: 2.5 mm; CoverWell Imaging Chambers PCI-A-2.5, Grace Bio-Labs, Bend, OR, USA), which was attached onto a glass slide. Approximately 30 paramecia were placed in the recording chamber, with a cover glass on top. The recording chamber was illuminated with a ring LED light (Model: LDR2-100SW2-LA, CCS Inc., Kyoto, Japan). The larva and paramecia were imaged with a stereoscope (SZX7, objective lens: DF PL 0.5X, Olympus, Tokyo, Japan) equipped with a CMOS camera (xiQ, Product number: MQ042RG-CM, Ximea, Lakewood, CO). Image acquisition was controlled by custom-made software developed on LabVIEW with the xiLIB (a LabVIEW Interface for XIMEA Cameras, https://decibel.ni.com/content/docs/DOC-29025). After time-lapse recordings, the number of paramecia left in the chamber was counted using the particle analysis function in ImageJ. Probability density function for the prey locations at the moment of prey capture behaviour was estimated using kernel density estimation in Matlab. Eye convergence was defined as the state in which the eye vergence angle^17, 50^ was more than 50°.

### Recording of the optokinetic response

A zebrafish larva was placed in 3% methylcellulose (3 mL) at the centre of a 35 ml-petri dish lid. The dish was placed in a custom-made motorised optokinetic drum (inner diameter 55 mm, height 80 mm). On the inner wall of the optokinetic drum, vertically striped sinusoidal gratings (360° /18 cycles) were present. The drum was rotated at a constant speed of 360° /7 s for the optokinetic stimulation.

### Ablation of the pretectal cells by a two-photon laser

For the behavioural study of the pretectum-ablated larvae, UAS:EGFP fish were used for ablation. Transgenic zebrafish larvae with UAS:EGFP;gSAIzGFFM119B on a *nacre* background were anaesthetised with tricaine throughout the laser ablation procedure. At 4 dpf, the larvae were mounted in 2% low-melting agarose. A two-photon laser scanning microscope (LSM7MP, Carl Zeiss, Germany) was used to ablate pretectal cells by a two-photon laser (at 880 nm, maximum power: approximately 2300 mW). Pretectal cells were observed with a 63X objective lens (W Plan-Apochromat 63x/1.0, Carl Zeiss, Germany) and images of 256 x 256 pixels were acquired, and irradiated at maximum power with the laser using the “bleaching” function at a scanning speed of 13.93 s / 256 pixels (134.42 μm) in the imaging software (ZEN 2011, Carl Zeiss). Bleaching was applied only on a small region (ROI) on each pretectal cell body to avoid any damage to the surrounding cells/tissue. Typically, about 10 cells were targeted on each focal plane, processing through 4 or 5 planes that covered the entire pretectal area labelled by the EGFP reporter. Because the efficiency of the ablation varied among cells, after going through the all focal plane, the cells were checked for the fluorescence. Fluorescent cells with insufficient ablation were subjected to laser irradiation again. One control larva (“No laser control”) and one larva for ablation were mounted in agarose in a dish for each ablation procedure and then recovered from the agarose. Prey capture ability in these larvae was tested in the following day (at 5 dpf). Effectiveness of ablation was confirmed after prey capture assay by fluorescent microscopy. Small number of fluorescent cells were usually found probably because of later onset of Gal4 gene expression in newly differentiated cells^51^. As another control, cells in the olfactory bulb UAS:EGFP;gSAIzGFFM119B larvae were laser-ablated in the same manner.

For the calcium imaging study of the pretectum-ablated larvae, UAShspzGCaMP6s, instead of UAS:EGFP, was used for ablation. GCaMP6s-expressing cells were irradiated by a two-photon laser at the wavelength of 800 nm. Ablation using GCaMP6s was performed 6~7 hours before behavioural assay on the same day at 5 dpf.

### Examination of Gal4 expression patterns

Transgenic zebrafish larvae with UAS:EGFP;gSAIzGFFM119B, UAS:RFP;gSAIzGFFM119B, UAS:EGFP;hspGFFDMC76A, or UAS:RFP;hspGFFDMC76A on a *nacre* background or with 1-phenyl 2-thiourea-treatment were anaesthetised using tricaine and observed under a Zeiss LSM7MP microscope using the Z-stack function.

### Labelling of the retinal ganglion cell axons

A lipophilic dye, 1, 1’-Dioctadecyl-3, 3, 3’,3’-tetramethylindocarbocyanine perchlorate (DiI, Cat # 468495, Sigma-Aldrich, St. Louis, MO, USA) was dissolved in N,N-Dimethylformamide (Cat # D4551, Sigma Chemicals, Balcatta, WA, USA) at 2 mg/mL. Transgenic zebrafish larvae harbouring UAS:EGFP;gSAIzGFFM119B on a *nacre* background were fixed in 4% paraformaldehyde for 1 h, embedded in 1% agarose covered with phosphate-buffered saline (PBS), injected with the DiI solution into the eye, and kept for 12 h to allow the DiI to diffuse along the retinal ganglion cell axons. The larvae were then mounted in 1.5% low-melting agarose and observed with a two-photon microscope (LSM7MP, Carl Zeiss, Germany) to obtain EGFP and DiI fluorescence images.

### Sparse cell labelling of the pretectal cells

A Tol2 vector DNA construct of UAS:EGFP (250 ng/nL; without transposase mRNA) was injected into the eggs of UAS:RFP;gSAIzGFFM119B at the 1-cell to 4-cell stages. Typically, we observed sparsely labelled pretectal cells at the larval stage in ~5% of the injected eggs. Single cell morphology of the sparsely labelled cells was examined under a two-photon microscope. The projection patterns of the neurites were analysed in IMARIS image processing software (Bitplane, South Windsor, CT, USA).

### Statistical tests

Statistical tests were performed in R (https://www.R-proiect.org/). A difference with p<0.05 was regarded as significant. Student’s t-test was used to compare two experimental groups of interest. Tukey’s HSD (honestly significant difference) test was used in multiple comparisons. Binomial test was used to assess the laterality in the neuronal response.

### Data availability

All data and codes used for the analysis are available from the authors upon request.

## Acknowledgements

We thank M. Suzuki, N. Mouri, and M. Mizushina for their help with the fish husbandry and A. Ito, F. Mizuno, Y. Kanebako and M. Iwasaki for their technical assistance. This study was supported by JSPS KAKENHI Grant Numbers JP25290009 and JP25650120, and also partly supported by JSPS KAKENHI Grant Numbers JP15H02370 and JP16H01651, and NBRP from Japan Agency for Medical Research and Development (AMED). This work was also supported in part by the Center for the Promotion of Integrated Sciences (CPIS) of SOKENDAI.

## Author contributions

AM and KK designed the experiments, and AM performed the experiments. PL, MI, GA, and KK generated the zebrafish Gal4 lines.PL and MI characterised the Gal4 expression in the adult brain. DA characterized and established the UAS:zBoTxBLCGFP transgenic line. AM and KK wrote the manuscript.

## Author information

The authors declare no competing financial interests. Requests for materials should be addressed to KK.

---

**Supplementary Movie 1. Activity of the inferior lobes of the hypothalamus (ILH) during prey capture behaviour (Supplement to Figure 1b-c)**

Calcium imaging of a 4 dpf UAS:GCaMP6s; hspGFFDMC76A larva with no prior feeding experience. Upper left: Raw fluorescent images after image registration to fix the head. Upper right: Image that was normalized to the frame before the Ca increase and is presented in pseudo-colours. Bottom: Raw fluorescent images with contrast enhanced to show the paramecium.

**Supplementary Movie 2. Hypothalamic response to prey in an agarose-embedded larva (Supplement to Figure 1d)**

Calcium imaging in a 5 dpf UAS:GCaMP6s; hspGFFDMC76A larva with a paramecium in the recording chamber. The trunk of the larva was embedded in agarose.

**Supplementary Movie 3. Pretectal response to prey in a 4 dpf zebrafish larva (Supplement to Figure 2b)**

Calcium imaging in a 4 dpf UAS:GCaMP6s;gSAzGFFM119B larva with a paramecium in the recording chamber.

**Supplementary Movie 4. Pretectal activity in the presence of a paramecium in a 6 dpf zebrafish larva embedded in agarose (Supplement to Figure 2d)**

Calcium imaging in a 6 dpf UAS:GCaMP6s;gSAzGFFM119B larva with a paramecium in the recording chamber. This movie is an example of the recordings in which eye convergence occurred. The movie is replayed in real time. Note that pretectal activation preceded the cerebellar activation that was associated with the eye movements. The mean intensities of the fluorescence in the regions of interest (ROIs) set on the left and right pretectal areas (green and magenta, respectively) and cerebellum (black and blue, respectively) are shown. The change in fluorescence intensity change (F/F0) is shown on the vertical axis (range: 0.8-2.5).

**Supplementary Movie 5. Pretectal activity in the presence of a paramecium in a 6 dpf larva embedded in agarose (no eye convergence) (Supplement to Figure 2d)**

A 6 dpf UAS:GCaMP6s;gSAzGFFM119B larva with a paramecium in the recording chamber. This movie is an example of the recordings in which no eye convergence occurred. The movie is replayed in real time. The mean fluorescence intensity in the ROIs set on the left and right pretectal areas (green and magenta, respectively) and cerebellum (black and blue, respectively) are shown. The change in fluorescence intensity (F/F0) is shown on the vertical axis (range: 0.8-2.5).

**Supplementary Movie 6. Pretectal response to a moving spot (Supplement to Figure 2e and f)**

A moving spot projected on a screen was shown to a 4-dpf UAS:GCaMP6s;gSAzGFFM119B zebrafish larva. The neuronal activity was recorded with a spinning-disk confocal microscope. A schematic of the visual stimulus is shown at the top (not to scale).

**Supplementary Movie 7. Prey capture is abolished in pretectum-ablated zebrafish (Supplement to Figure 2m)**

The gSAzGFFM119B-labelled pretectal cells were bilaterally ablated with a two-photon laser at 4dpf, and the zebrafish larvae were behaviourally tested at 5dpf. Left: a control larva that was embedded in agarose and kept in the same dish as the ablated larva. Right: a larva that was subjected to bilateral ablation of the pretectal cells by a two-photon laser. Duration of the movie: 11 min. Note that the paramecia, which appear as yellowish dots, are consumed in the control chamber but not in the pretectum-ablated chamber.

**Supplementary Movie 8. gSAIzGFFM119B-labelled pretectal cells project to the inferior lobe of the hypothalamus (Supplement to Figure 3a)**

Two-photon confocal images (z-stack) of a 5 dpf UAS:EGFP;gSAIzGFFM119B zebrafish larva. This Gal4 line labels the forebrain, a nucleus in the pretectal area, preglomerular nuclei, and cerebellum, which is not included in the focal planes shown in this movie. The focal plane moves ventrally from the pretectum to the hypothalamus. The axons of the pretectal cells are indicated by an arrowheads.

**Supplementary Movie 9. Correlated calcium signals in the pretectal areas and the inferior lobes of the hypothalamus in response to prey in an agarose-embedded larva (Supplement to Figure 3b)**

Calcium imaging in a 5 dpf UAS:GCaMP6s;gSAzGFFM119B;hspGFFDMC76A larva with a paramecium in the recording chamber. The trunk of the larva was embedded in agarose. F/F0 is shown in pseudo-colours.

**Supplementary Figure 10. gSAIzGFFM119B-labelled pretectal cells project to the precerebellar area as well as to the inferior lobes of the hypothalamus (ILH)**

A 5 dpf gSAIzGFFM119B;UAS:RFP larva was observed with a spinning-disk confocal microscope. The larva was tilted by approximately 17° (tail side up relative to the head) so that the pretectal projections to the cerebellar area were in the horizontal plane (for better spatial resolution of the image).

**Supplementary Figure 1.**
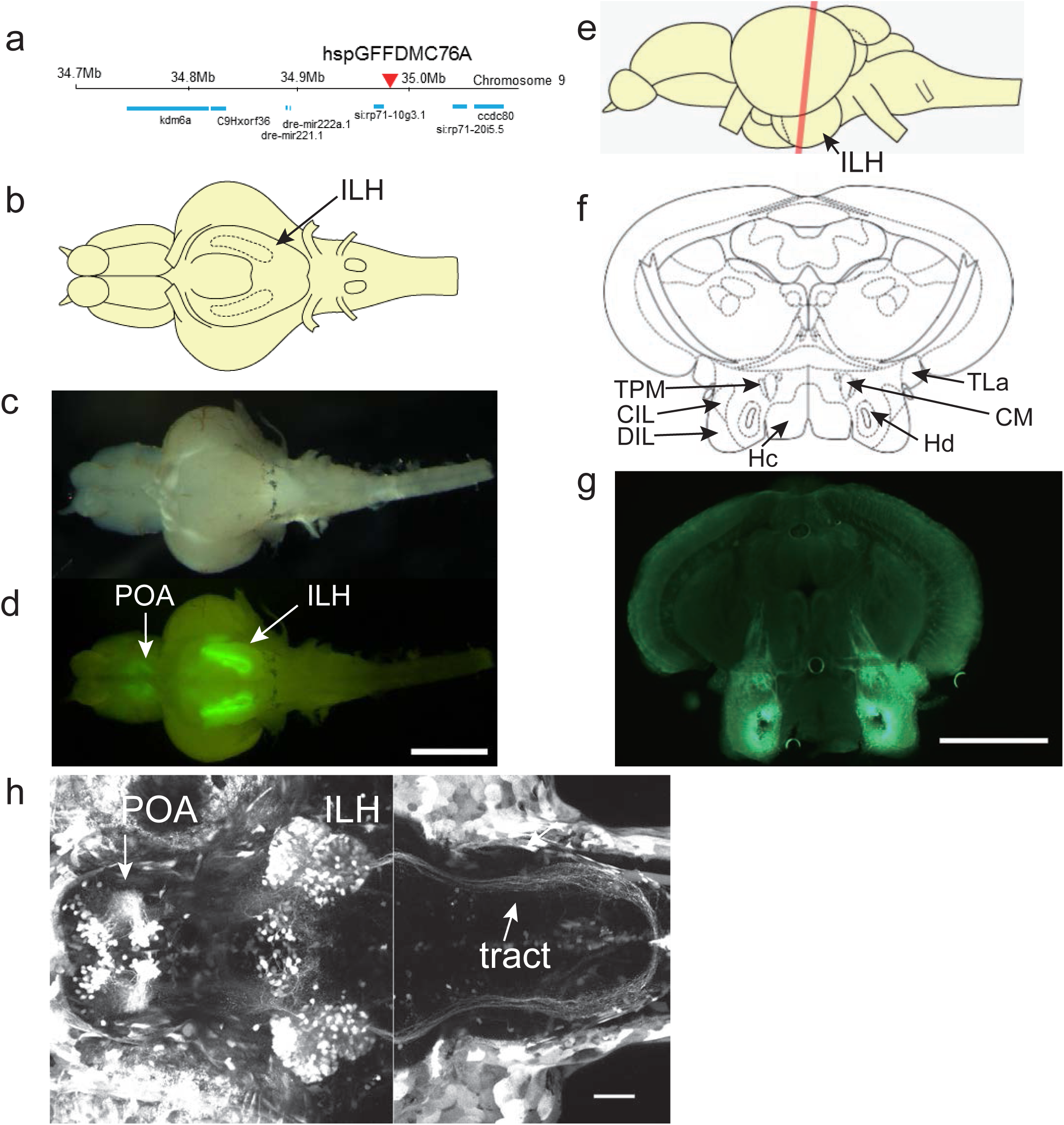
UAS:EGFP reporter gene expression in the hspGFFDMC76A Gal4 line. **a,** Insertion site of the hspGFFDMC76A was identified in an intergenic region on chromosome 9. **b.** Schematic of a ventral view of the brain of an adult zebrafish. **c,** Ventral view of a dissected brain of an adult UAS:EGFP;hspGFFDMC76A fish. **d,** EGFP fluorescence in the brain shown in **b**. Scale bar: 1 mm. **e,** Schematic side view of the adult brain. The red line indicates the positions of the sections in **f** and **g. f,** Annotations based on Wulliman et al., 1996. **g,** EGFP fluorescence of a coronal section of the brain of an adult UAS:EGFP;hspGFFDMC76A zebrafish. Scale bar: 0.5 mm. **h,** UAS:EGFP expression in the hspGFFDMC76A Gal4 line at 5 dpf. Projected z-stack images obtained by two-photon laser microscopy. Left: z-stack projection of 155 slices (1 pm-step). Right: z-stack projection of 135 slices (1 pm-step). The positions of the focal planes of the two z-stacks overlap by 35 um (the right one more dorsal) along z-axis. Scale bar: 50 pm. CIL, central nucleus of the inferior lobe; CM, corpus mamillare; DIL, diffuse nucleus of the inferior lobe; Hc, caudal zone of periventricular hypothalamus; Hd, dorsal zone of periventricular hypothalamus; ILH, inferior lobes of the hypothalamus; POA, preoptic area; TLa, torus lateralis; TPM, tractus pretectomamillaris.

**Supplementary Figure 2.**
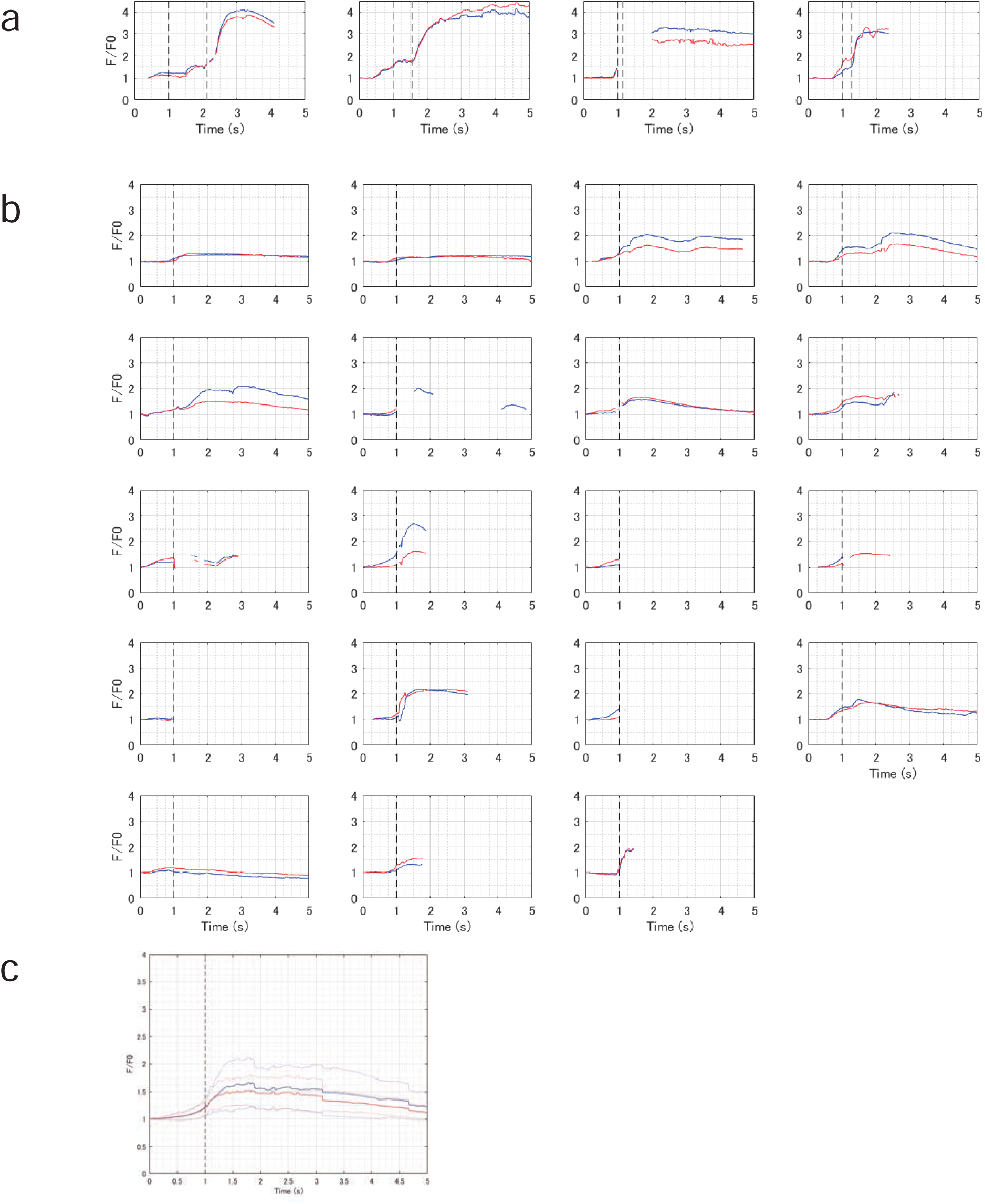
Neuronal activity in the ILH in free-swimming larvae. **a,** Calcium signals in the left (blue) and right (red) ILH during prey capture behaviour in 4 dpf UAShspGCaMP6s;hspGFFDMC76A larvae. Four examples of successful prey capture events that were observed in 4 larvae (example on far left is also shown in Figure 1c). The black dotted vertical line represents the time of eye convergence. The grey dotted vertical line represents the time of the completion of the prey capture. The gaps in the graph indicate times when the image registration failed due to large/rapid movements of the larvae. **b,** Calcium signals in the ILH during prey capture behaviour that were aborted before capture (19 examples in 4 larvae). The black dotted vertical line represents the time of eye convergence. **c,** Average calcium signals shown in **b**. The graphs are aligned relative to the timing of eye convergence. The steps in the graph are artefacts from missing data. Note that the calcium signals starts to increase before eye convergence. The solid lines represent the mean, and the dotted lines represent the S.D.

**Supplementary Figure 3.**
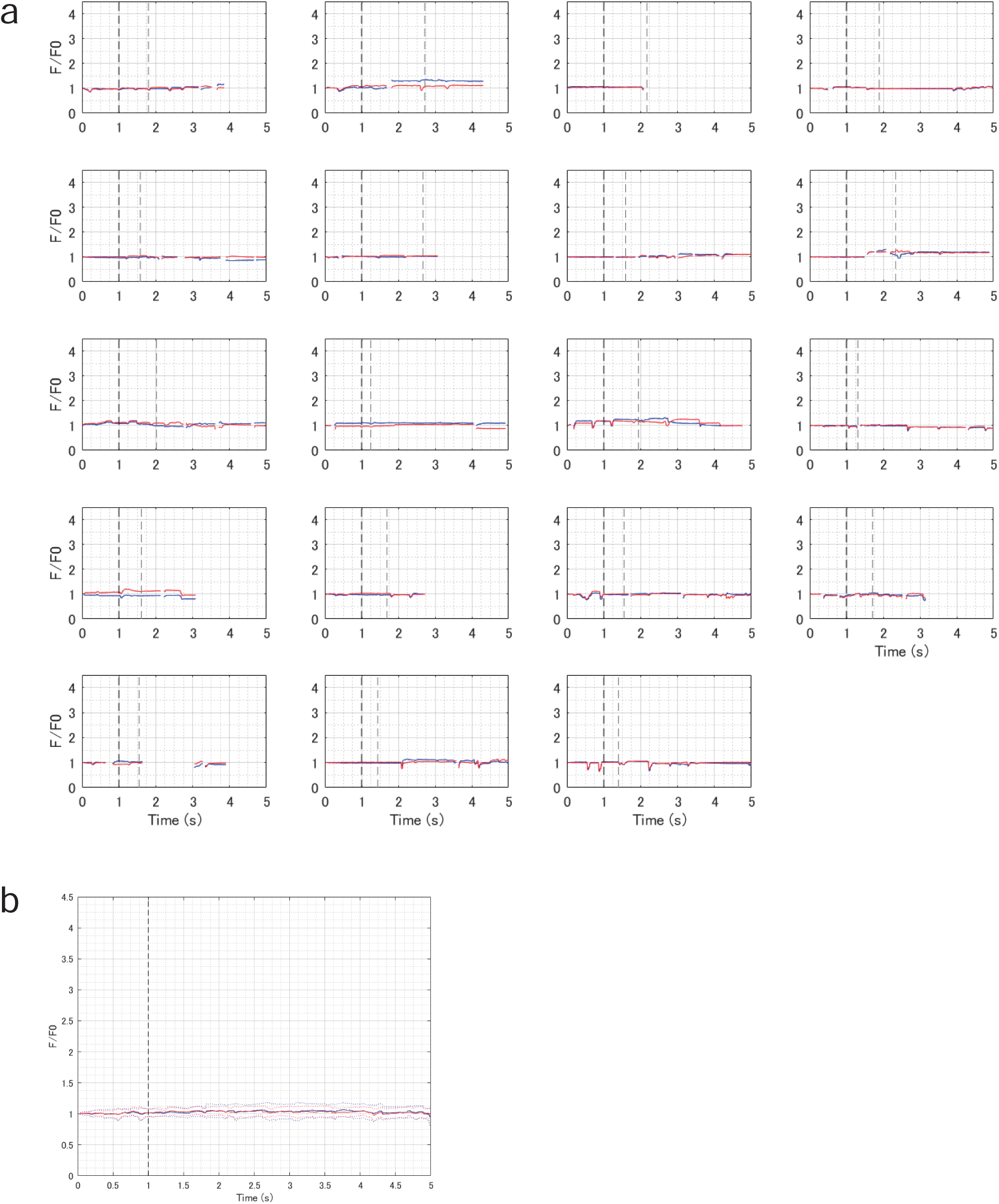
EGFP fluorescence imaging in the ILH in free-swimming larvae. **a,** EGFP fluorescence signals in the left (blue) and right (red) ILH during prey capture behaviour in 5 dpf UAS:EGFP;hspGFFDMC76A larvae. Nineteen examples of successful prey capture events were observed in 3 larvae. The black dotted vertical line represents the time of eye convergence. The grey dotted vertical line represents the time of prey capture. The gaps in the graph indicate times when the image registration failed due to large/rapid movements of the larvae. **b,** Averaged EGFP fluorescence signals shown in **a**. The graphs are aligned relative to the timing of eye convergence. The solid lines represents the mean, and the dotted lines represent the S.D.

**Supplementary Figure 4.**
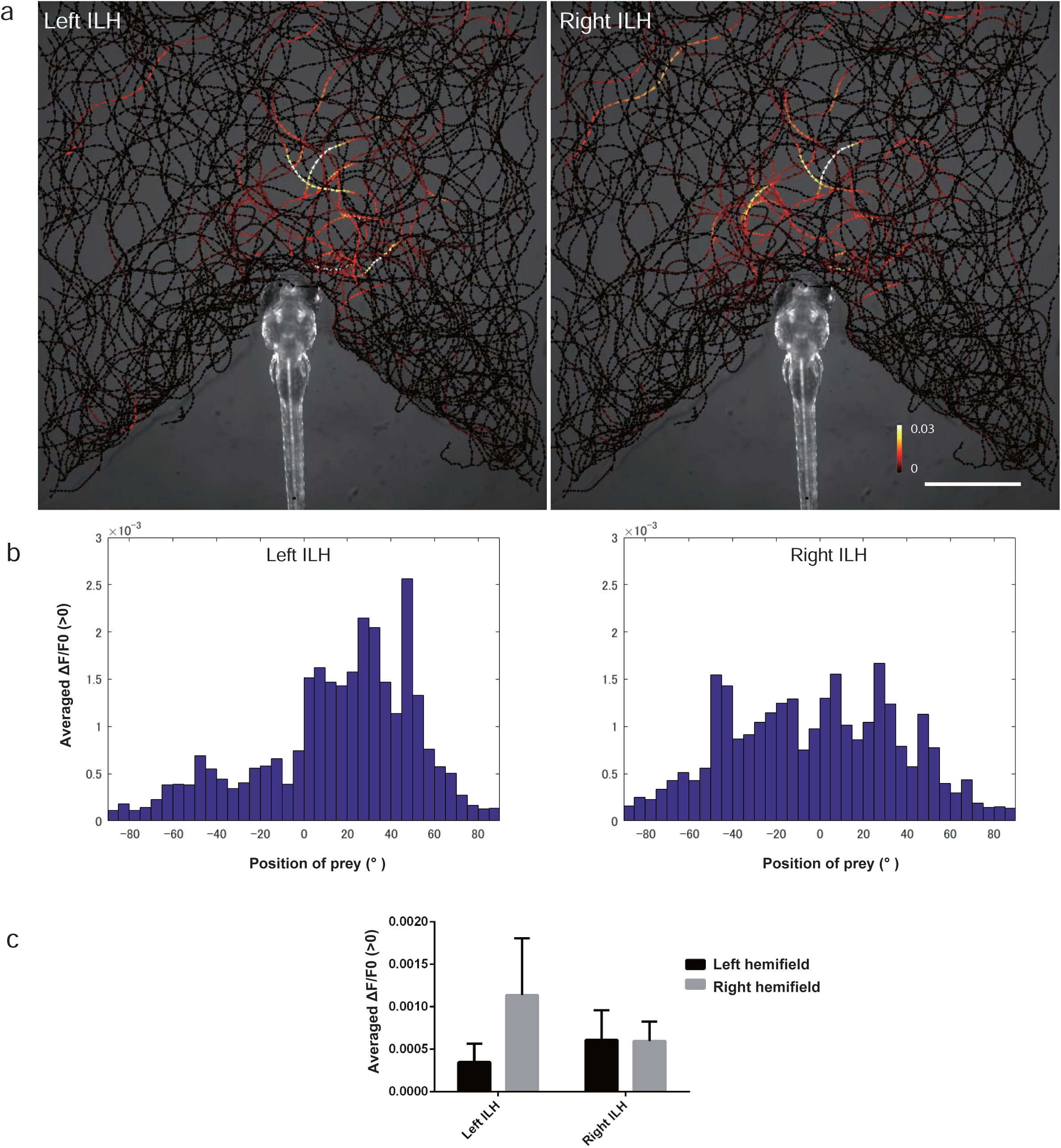
Activity in the ILH at the sight of prey. **a,** The trajectories of single paramecia over 942 s are shown with the colour-coded changes in the intensity of GCaMP6s fluorescence in the left and right ILH areas in 5 dpf UAShspGCaMP6s;hspGFFDMC76A larvae. The data from 4 larvae were merged into a single larval image. Scale bar: 1 mm. The length of the arrowhead indicates the distance travelled by the paramecium in 60 ms. **b,** Average increase in the Ca signals in each 5° bin. The data are the same as those shown in **a**. **c,** Average increase in the Ca signals in the left and right hemifields. The data are the same as those shown in **a**. Mean ± S.D.

**Supplementary Figure 5.**
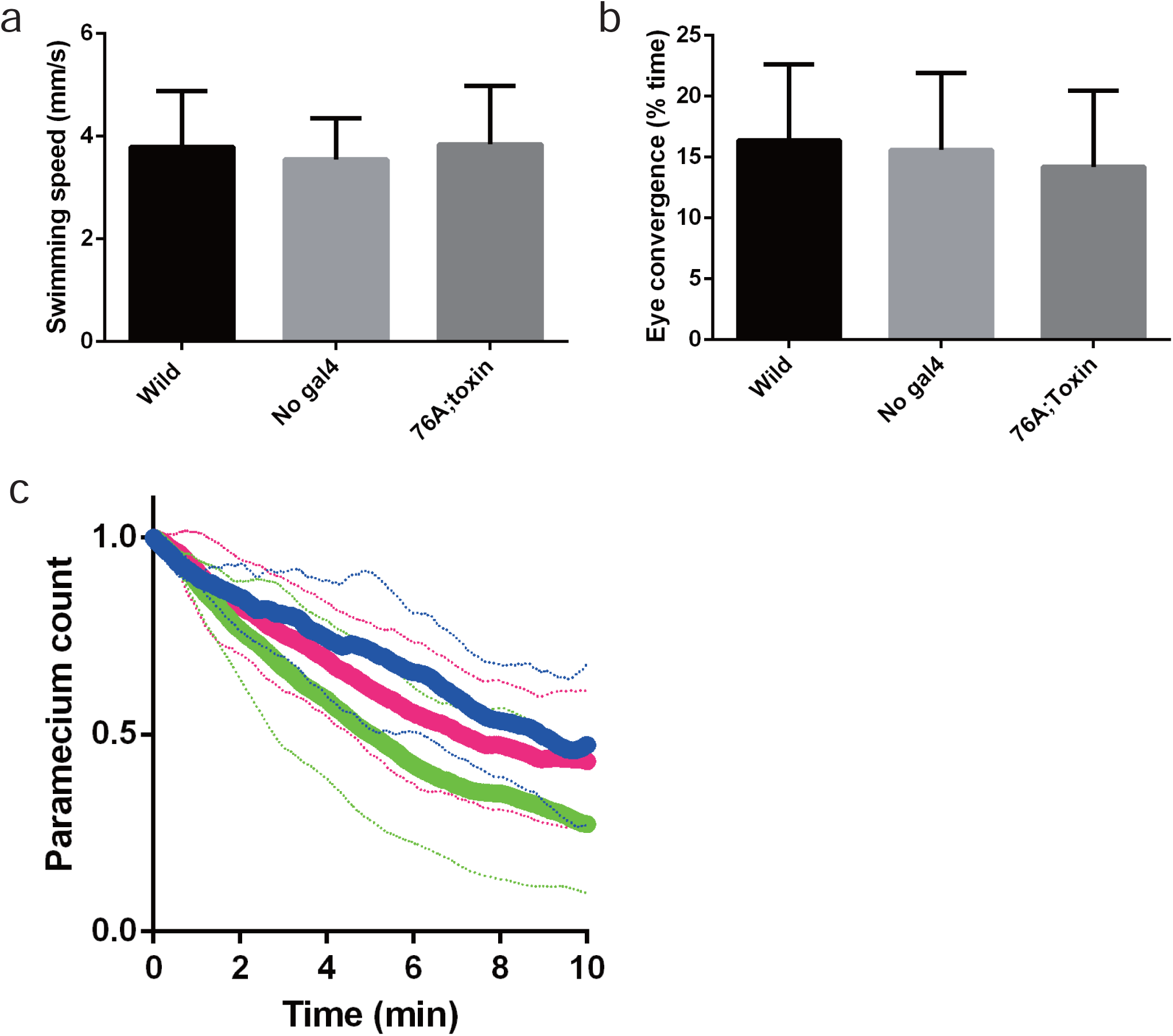
Effects of botulinum toxin expression. **a,** Swimming speeds of wild-type (n=13), UAS:zBoTxBLCGFP (n=18), and UAS:zBoTxBLCGFP; hspGFFDMC76A larvae (n=6). Mean ± S.D. **b,** Eye convergence during 10 min with prey in wild-type (n=13), UAS:zBoTxBLCGFP (n=18), and UAS:zBoTxBLCGFP; hspGFFDMC76A larvae (n=6). Mean ± S.D. **c,** Paramecium consumption in 10 min in UAS:zBoTxBLCGFP;UAS:EGFP; SAGFF(LF)27A (magenta, n=9), UAS:zBoTxBLCGFP (blue, n=5) and UAS:EGFP;SAGFF(LF)27A (green, n=9) larvae. The solid lines represent the mean and the dotted lines represent the S.D. Two-tailed t-test between UAS:zBoTxBLCGFP and UAS:EGFP;SAGFF(LF)27A groups, p=0.07.

**Supplementary Figure 6.**
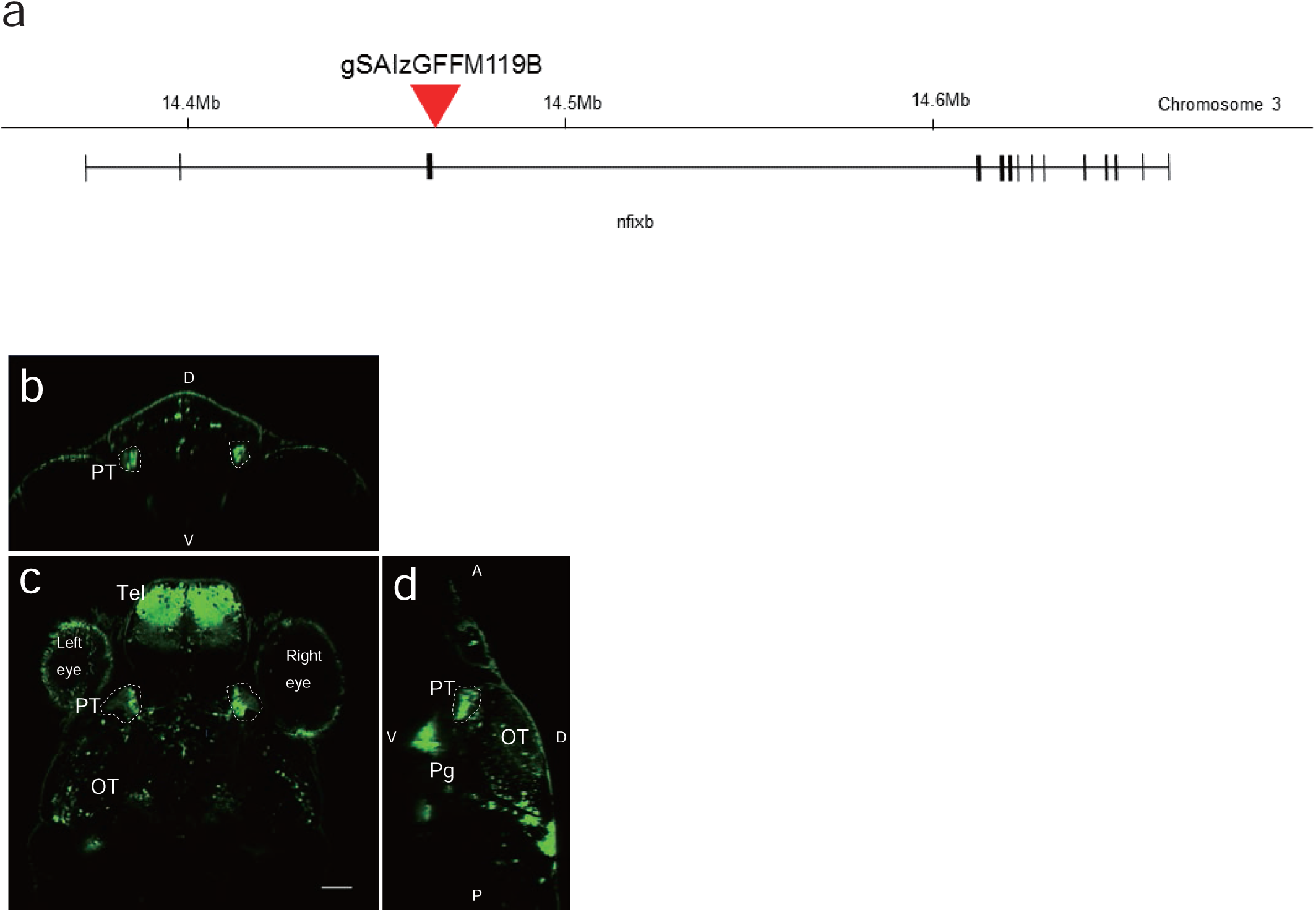
The Gal4 insertion site in the gSAIzGFFM119B gene-trap Gal4 line and UAS:EGFP reporter gene expression. **a,** Insertion site of the Gal4 construct in the gSAIzGFFM119B line. The insertion site was identified within the *nfixb* gene on chromosome 3. **b-d,** Confocal images of the expression of the UAG:EGFP reporter, driven by Gal4 in the gSAzGFFM119B line. **b,** Coronal optical section. **c,** Dorsal view of a single optical section showing the positions of the pretectal cells. Scale bar: 50 μm. **d,** Sagittal view. D, dorsal; V, ventral; PT, pretectum; Tel, telencephalon; OT, optic tectum; Pg, preglomerular nuclei.

**Supplementary Figure 7.**
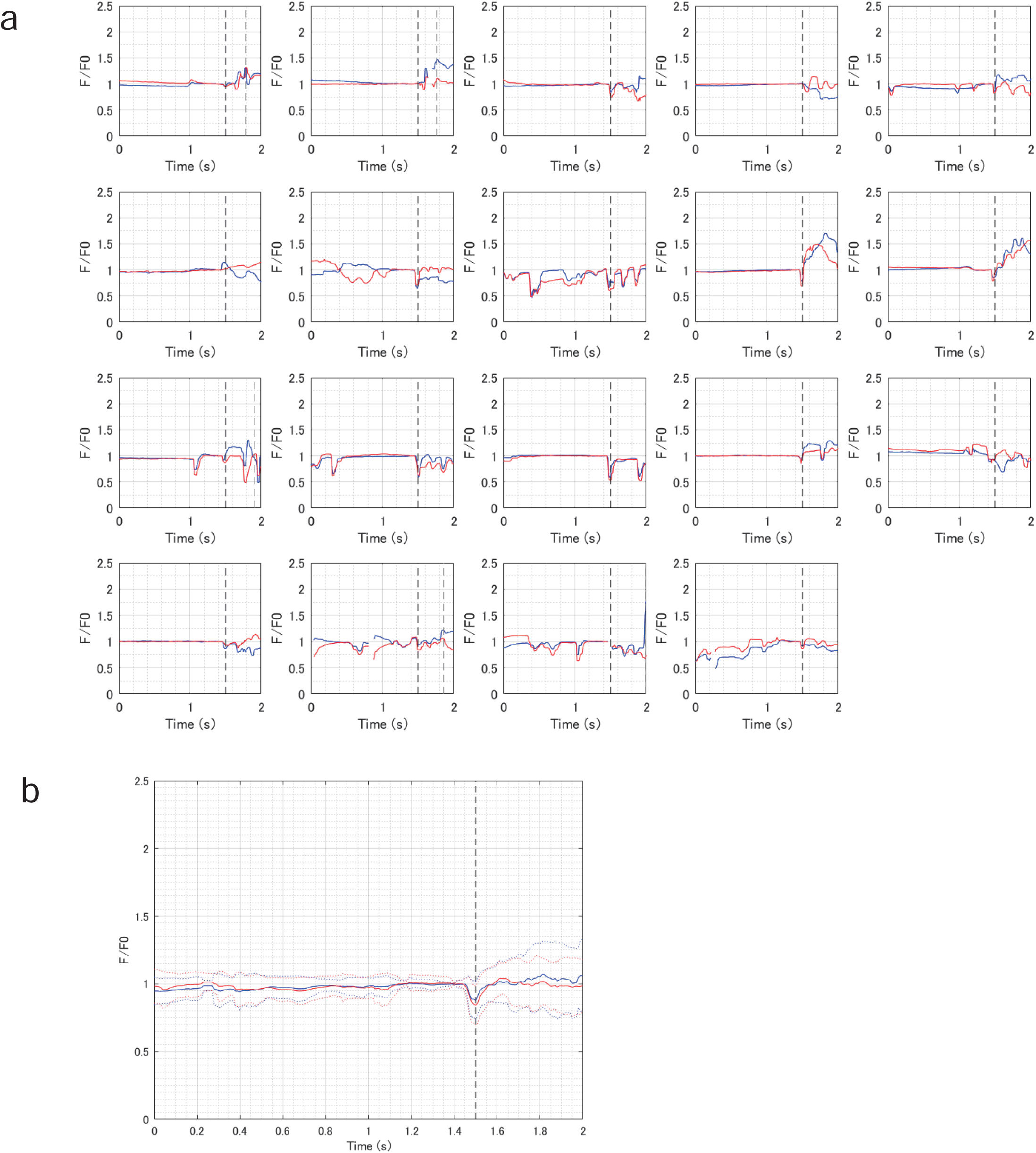
EGFP fluorescence imaging in the pretectum in free-swimming larvae. a, EGFP fluorescence in the left (blue) and right (red) pretectum during prey capture behaviour in 5 dpf UAS:EGFP; gSAzGFFM119B larvae. The 19 examples of successful prey capture events that were observed in 3 larvae are shown. The black dotted vertical line represents the time of eye convergence. The grey dotted vertical line represents the time of prey capture (in some graphs, it occurred outside the time range). The gaps in the graph indicate times when the image registration failed due to large/rapid movements of the larvae. **b,** Averaged EGFP fluorescence signals shown in **a**. The graphs are aligned relative to the timing of eye convergence. The solid lines represent the mean, and the dotted lines represent the S.D.

**Supplementary Figure 8.**
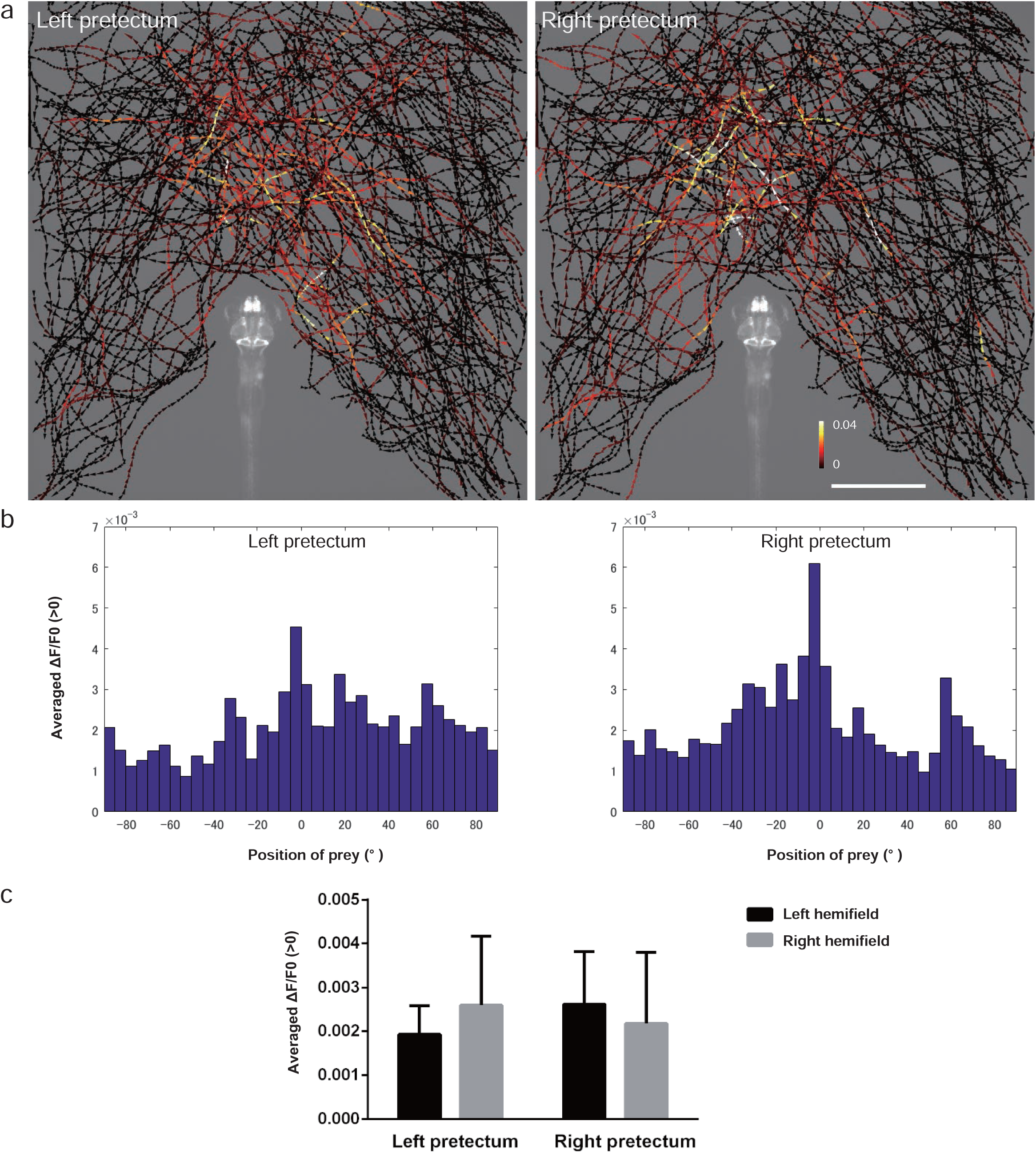
Activity in the pretectal cells at the sight of prey. **a,** The trajectories of single paramecia over 775 s are shown with the colour-coded changes in the intensity of GCaMP6s fluorescence in the pretectum in 5 dpf UAShspGCaMP6s; gSAIzGFFM119B larvae. The data from 4 larvae are merged into a single larval image. Scale bar: 1 mm. The length of the arrowhead represents the distance travelled by the paramecium in 60 ms. **b,** Average increase in the Ca signals in each 5° bin. The data are the same as those shown in **a**. **c,** Averaged increase in the Ca signals in the left and right hemifields. The data are the same as those shown in **a**. Mean ± S.D.

**Supplementary Figure 9.**
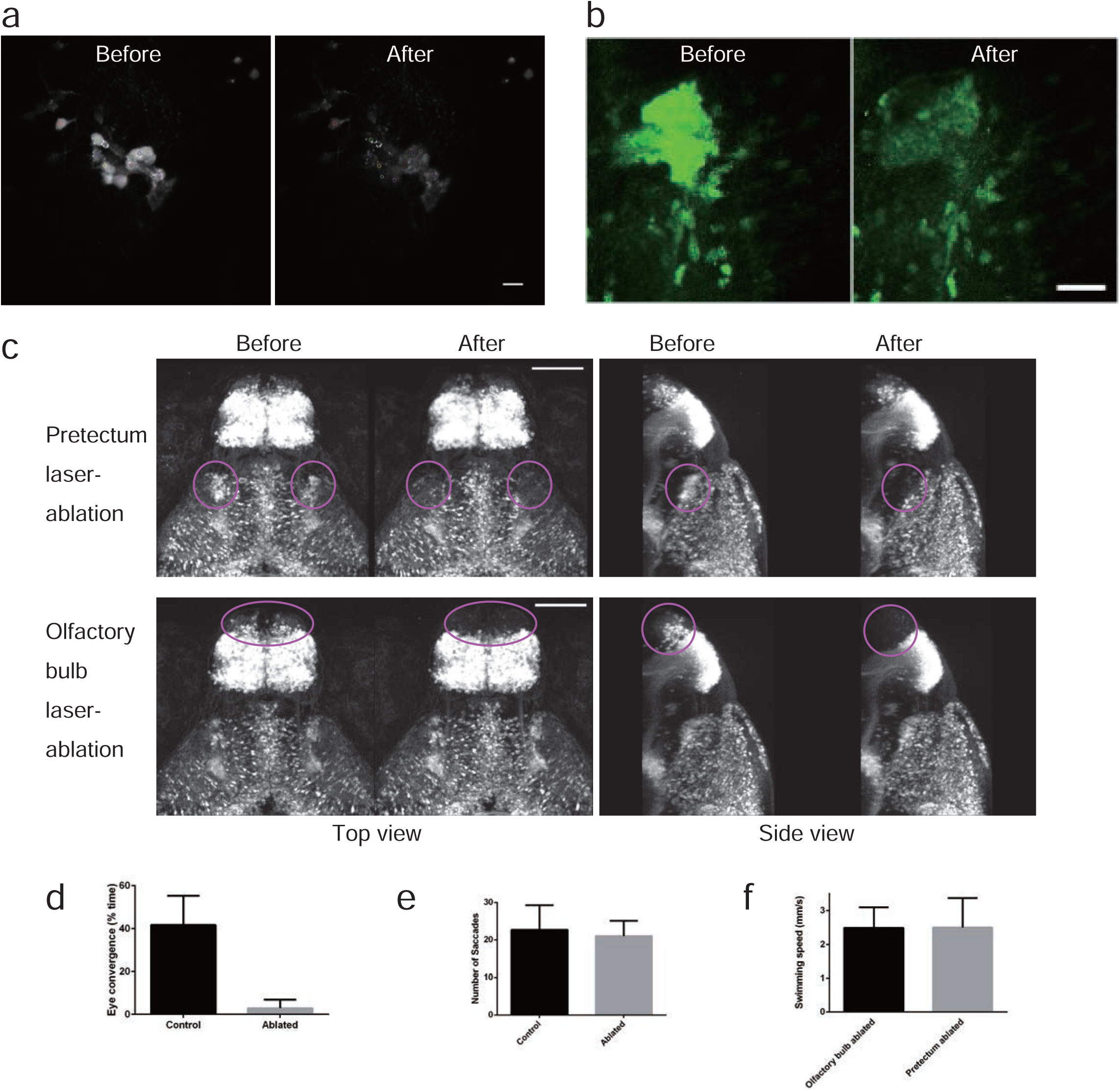
Ablation of the pretectal cells in UAS:EGFP;gSAIzGFFM119B larvae. **a,** Images of the EGFP fluorescence of the pretectal cells on a focal plane before and after two-photon laser ablation in a 4 dpf UAS:EGFP;gSAIzGFFM119B larva. The targeted cells are labelled with small coloured circles. Scale bar: 10 μm. **b,** Image of the EGFP fluorescence of the pretectal cells in a UAS:EGFP;gSAIzGFFM119B larva. z-stack images that covered the entire gregion of the 119B-labelled pretectal cells were projected onto one image. Before (left) and after (right) laser ablation. Scale bar: 25 pm. **c,** Absence of fluorescent cells after laser ablation. Top panel: pretectum ablation (encircled in magenta). Bottom panel: olfactory bulb ablation (encircled in magenta) as a control experiment. Scale bar: 100 pm. **d,** Eye convergence (indicator of prey capture, mean ± S.D.; Control, n = 3 untreated larvae; Ablated, n = 5 pretectum-ablated larvae; two-tailed t-test, p = 0.0008). **e,** Optokinetic response (mean ± S.D.; Control, n = 8 untreated larvae; Ablated, n = 6 pretectum-ablated larvae; two-tailed t-test p = 0.5920). **f,** Locomotor activity in olfactory bulb-ablated larvae (n=11) and pretectum-ablated larvae (n=10). The average swimming speeds in 10 min recordings are shown with S.D.

**Supplementary Figure 10.**
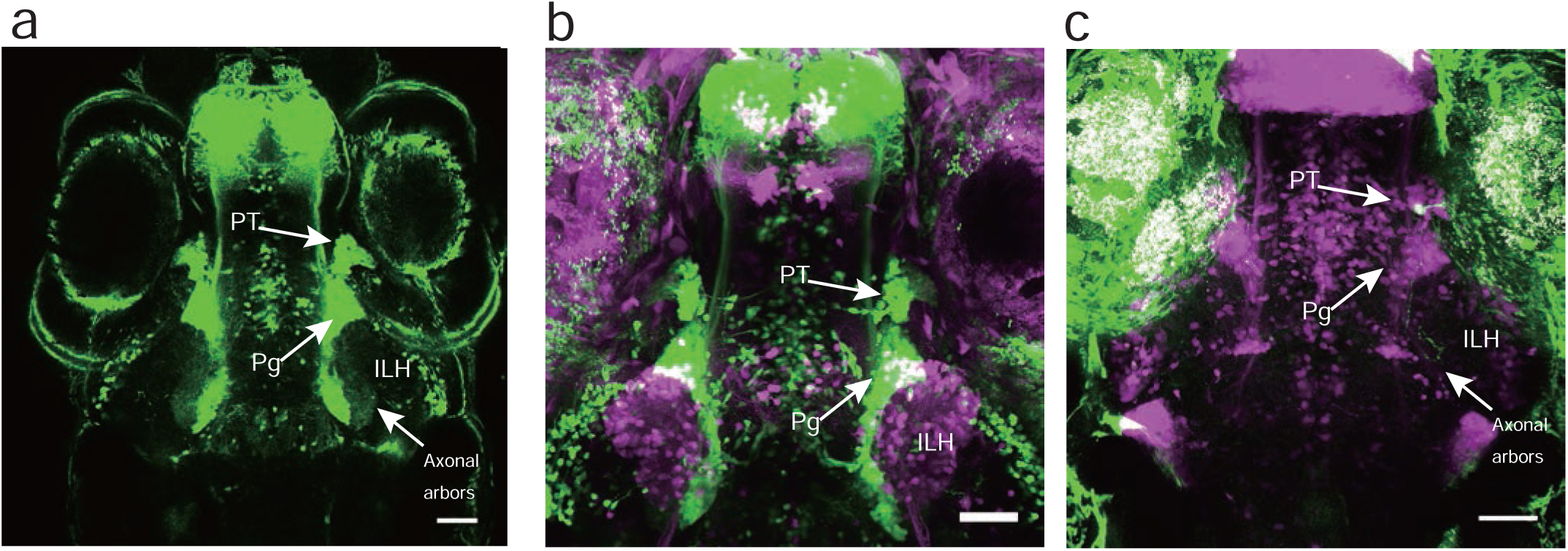
Projection of the 119B-pretectal cells to the ILH. **a,** Merged image of 3 optical sections (66.7 pm and 22 pm apart on the Z-axis in the ventral direction) depicting the pretectal area (PT), the preglomerular nuclei (Pg), and the axonal arbours of the pretectal cells that reached the ILH at 5 dpf. Scale bar: 50 μm. **b,** Merged image of a UAS:EGFP;gSAIzGFFM119B larva (green) and a UAS:EGFP;hspGFFDMC76A larva (magenta, a different individual), which shows the location of the ILH relative to the PT and Pg. Scale bar: 50 pm. **c,** A single pretectal (PT) cell labelled by a UAS:EGFP DNA injection (green) into a UAS:RFP;gSAIzGFFM119B background zebrafish (magenta) at the 1-cell stage and observed at 4 dpf. An axon is extending out of the soma of the labelled cell in the pretectal area and is projecting to the caudomedial ILH. Scale bar: 50 μm.

**Supplementary Figure 11.**
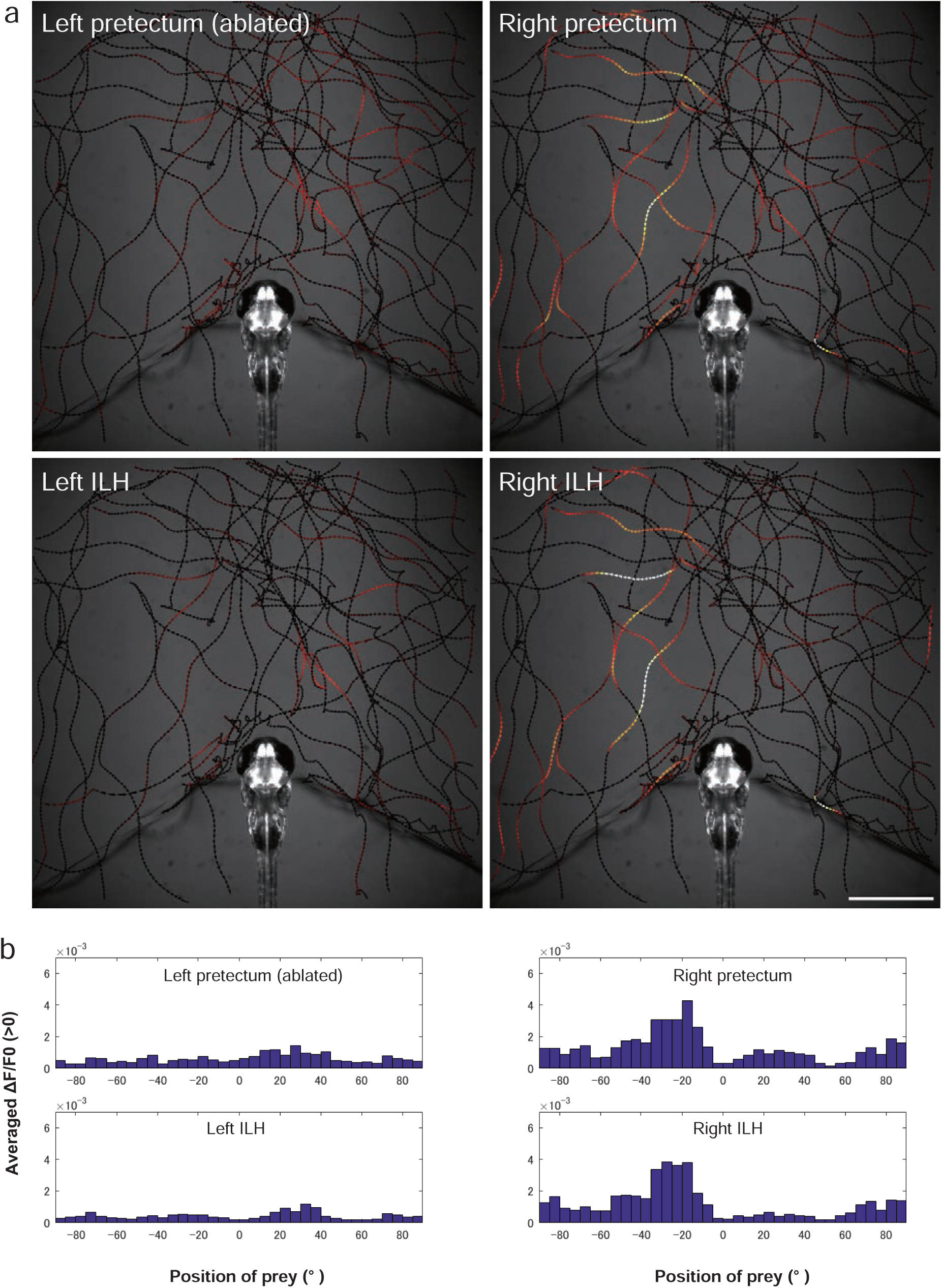
Activity in the pretectum and ILH at the sight of prey in a larvae subjected to unilateral pretectum ablation. **a,** Another example of larvae in the unilateral ablation experiments shown in Figure 3c. The trajectories of a single paramecium over 250 s are shown with the colour-coded changes in the intensity of GCaMP6s fluorescence in a 5 dpf UAShspGCaMP6s; hspGFFDMC76A; gSAIzGFFM119B larva that was subjected to laser ablation of the left pretectum. Scale bar: 1 mm. The length of the arrowhead represents the distance travelled by the paramecium in 60 ms. **b,** Average increase in the Ca signals in each 5° bin. The data are the same as those shown in **a**.

**Supplementary Figure 12.**
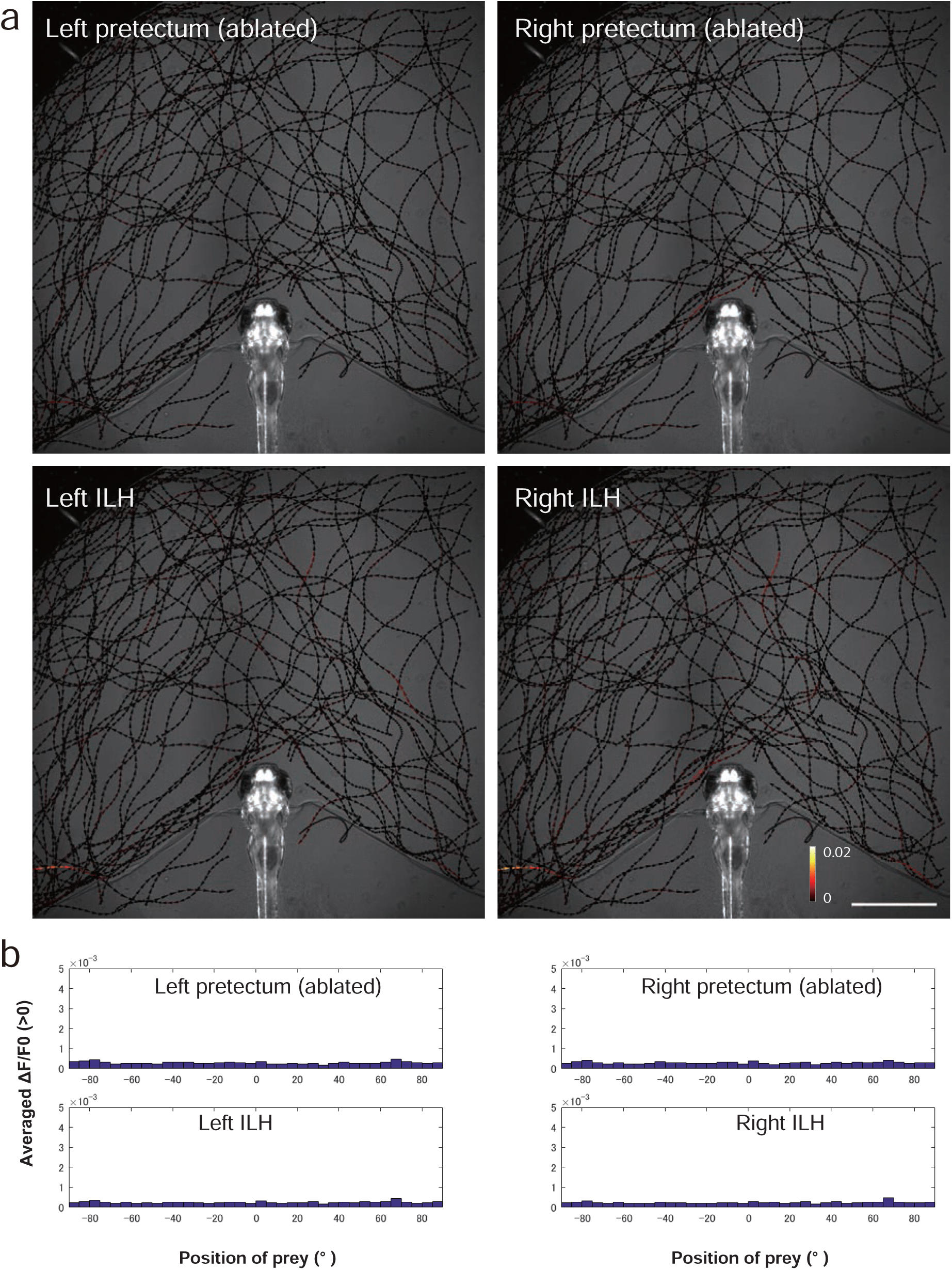
Activity in the pretectum and ILH at the sight of prey in a larvae subjected to bilateral pretecutm ablation. **a,** A representative example of the larvae in the bilateral ablation experiments. The trajectories of single paramecia over 324 s are shown with the colour-coded changes in the intensity of GCaMP6s fluorescence in a 5 dpf UAShspGCaMP6s;hspGFFDMC76A; gSAIzGFFM119B larva that was subjected to laser ablation of the bilateral pretectum. Scale bar: 1 mm. The length of the arrowhead represents the distance travelled by the paramecium in 60 ms. **b,** Average increase in the Ca signals in each 5° bin. The data are the same as those shown in **a**.

**Supplementary Figure 13.**
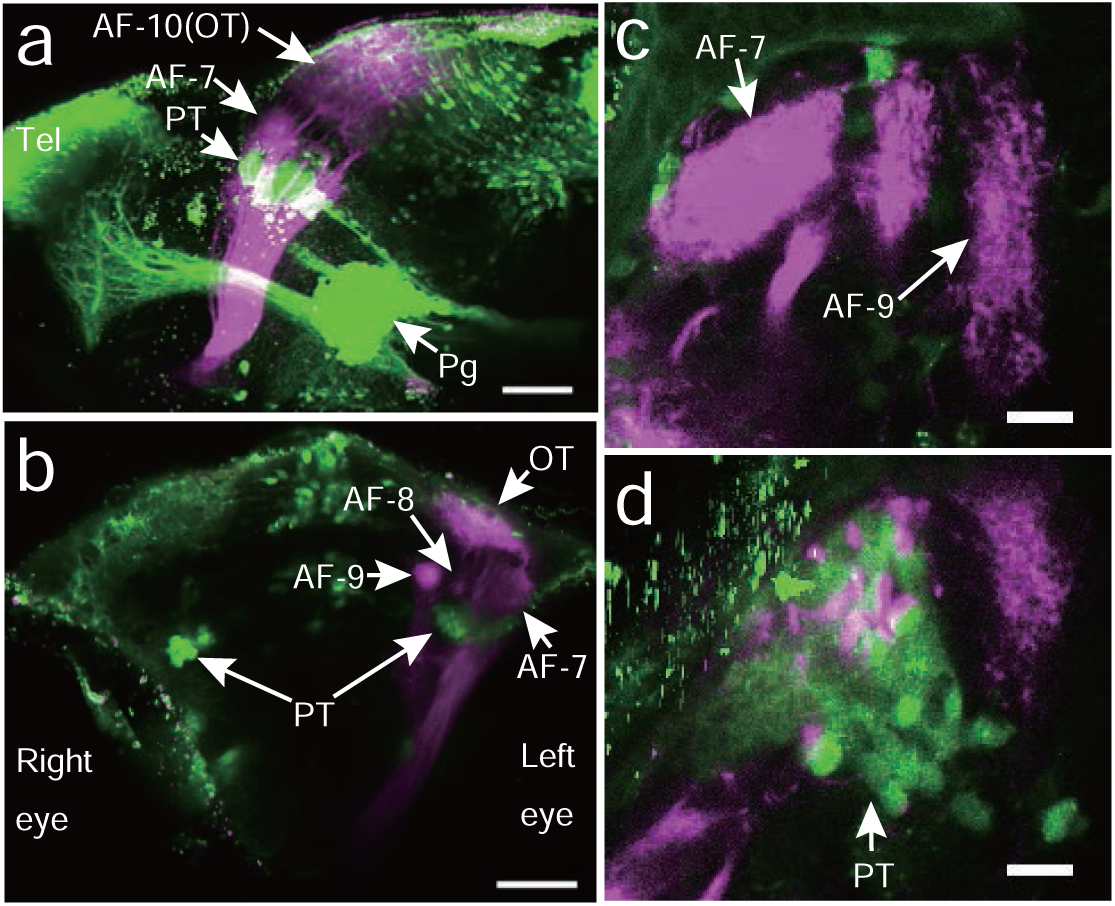
Locations of the 119B-pretectal area and the retinal ganglion cell arbours. Locations of the axonal arbourisation fields (AFs) of the ganglion cells (magenta) relative to the gSAzGFFM119B-labelled cells (green). DiI was injected into the right eye, and the left eye was removed for observation. **a,** Lateral view from the left side. Scale bar: 50 μm. **b,** Front view. Scale bar: 50 pm. **c,** Top view (focused on the AF-7 area). Scale bar: 5 μm. **d,** Top view (focused 26 μm below the AF-7 area). Scale bar: 5 μm. PT, pretectal area; OT, optic tectum; Tel, telencephalon; Pg, preglomerular nuclei; D, dorsal; V, ventral; A, anterior; P, posterior.

**Supplementary Figure 14.**
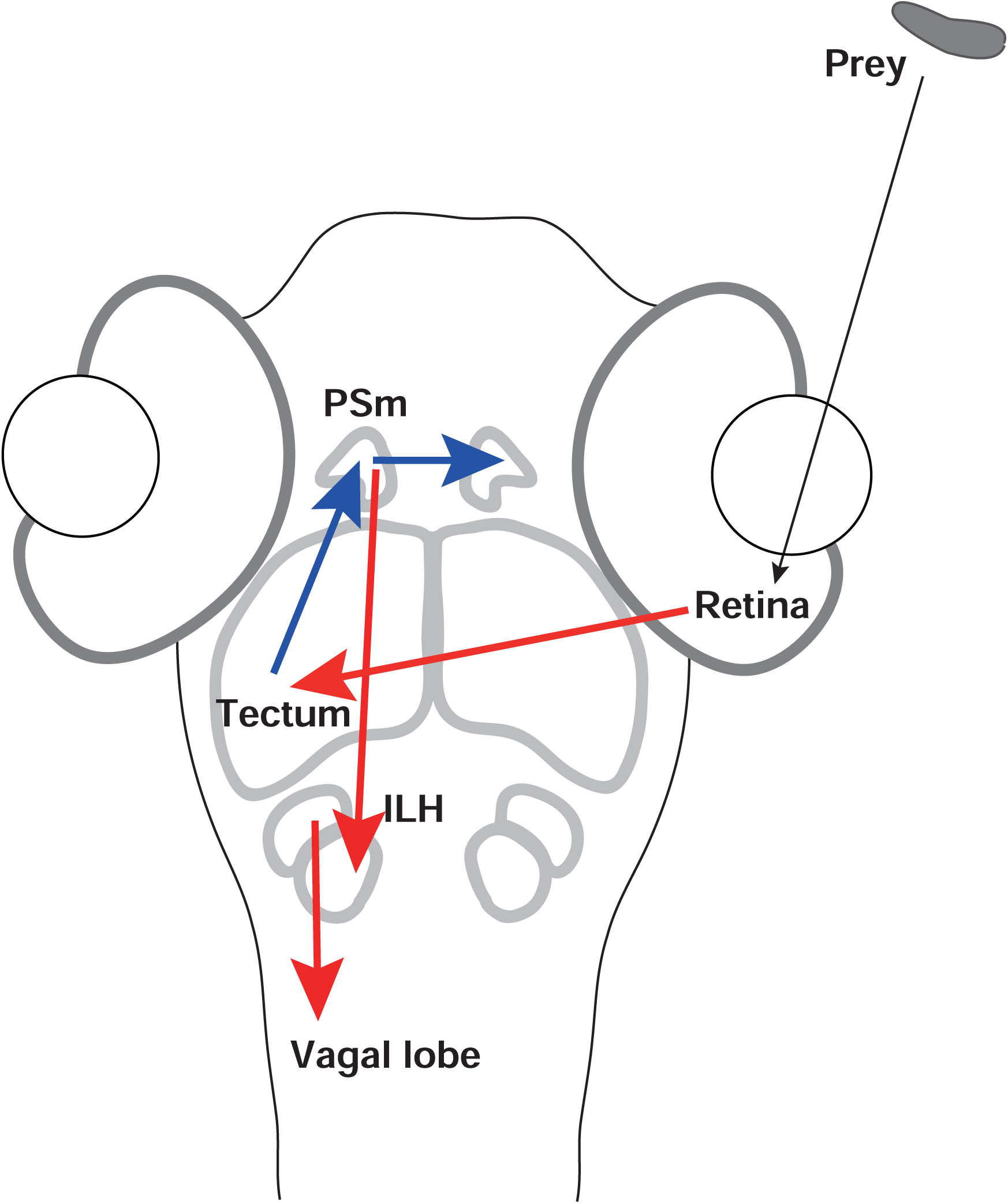
Model of the neural circuits of prey capture. The sight of prey forms an image that is projected onto the retina. The majority of the retinal ganglion cells project to the optic tectum of the midbrain. The nucleus pretectalis superficialis pars magnocellularis (PSm)(gSAIzGFFM119B-labelled cells) presumably receives inputs from the optic tectum and then projects to the posterior part of the inferior lobes of the hypothalamus (ILH) (hspGFFDMC76A-labelled cells). The anterior part of the ILH projects through the dorsal area of the hindbrain (presumably, vagal lobe). Bilateral activation of the pretectum might be produced by the interconnection between the left and right pretectal areas. The red arrows indicate the axonal projections that are based on the morphological data shown in this study. The blue arrows indicate speculative connections. The projections are shown only on the left side of the schematic for clarity.

